# Diversity and Divergence: Evolution of defense chemistry in the tropical tree genus *Inga*

**DOI:** 10.1101/2021.12.17.473194

**Authors:** Dale L. Forrister, María-José Endara, Abrianna J. Soule, Gordon C. Younkin, Anthony G. Mills, John Lokvam, Kyle G. Dexter, R. Toby Pennington, Catherine A. Kidner, James A. Nicholls, Oriane Loiseau, Thomas A. Kursar, Phyllis D. Coley

## Abstract

- Plants are widely recognized as chemical factories, with each species producing dozens to hundreds of unique secondary metabolites. These compounds shape the interactions between plants and their natural enemies. Here we explore how plants generate chemical diversity, and what evolutionary processes have led to novel compounds and unique chemical profiles.
- We comprehensively characterized the chemical profile of one-third of the species of tropical rainforest trees in the genus *Inga* (∼ 100, Fabaceae) and applied phylogenetic comparative methods to understand the mode of chemical defense evolution.
- We show that: 1) Each *Inga* species produces exceptionally high levels of phytochemical diversity, despite costs, tradeoffs and biosynthetic constraints. 2) Closely related species have highly divergent defense profiles, with individual compounds, major compound classes and complete profiles showing little to no phylogenetic signal. 3) We show that the evolution of a species’ chemical profile shows a signature of divergent adaptation, implying that it is advantageous for a species to have distinct chemistry from close relatives to avoid shared natural enemies. 4) Finally, we hypothesize a model where deep homology of biosynthetic pathways and rapid changes in regulatory mechanisms may better explain the observed large shifts in defense chemicals between closely related taxa.

## Introduction

For sessile organisms like plants, secondary metabolism plays a fundamental role in mediating biotic interactions ranging from mutualisms (e.g. pollination) to antagonisms (e.g. competition and defense). Plant secondary metabolites (PSMs, sometimes referred to as specialized metabolites), which are classically considered nonessential for basic cellular function, are exceedingly diverse, with nearly 1,000,000 predicted to exist across the plant kingdom (1). It has long been thought that this incredible diversity strongly influences the structure of plant-herbivore interactions (2–4). The evolution of novel compounds or unique combinations of compounds (hereafter, chemical profile) can be highly adaptive, increase plant fitness, and facilitate coexistence between related species (5–7). Thus, understanding the origin and maintenance of PSM diversity is central to understanding both the evolution and ecology of plant-herbivore interactions.

Much of the theoretical and empirical literature supports the idea that selection has placed a premium on chemical diversity in plants (8–13). A species’ chemical profile is thought to arise from a diverse set of selective pressures exerted by a multitude of herbivores, pathogens, and mutualists (8, 12). For example, increased phytochemical diversity in tropical forests decreases the number of herbivore species associated with a given host (12, 14) and can reduce rates of herbivory (15). In addition to producing a diverse set of compounds, recent studies have highlighted the importance for a given species to maintain a unique chemical profile relative to other species in its community (7, 16, 17). While there is a clear consensus on the value of phytochemical diversity, the underlying evolutionary processes that generate chemical diversity in plant lineages remain widely debated (11).

Despite the apparent adaptive advantages of phytochemical diversity, metabolic and biosynthetic constraints may limit a species’ ability to produce a diverse array of compounds. Investment in chemical defense is expensive both in terms of the carbon and nitrogen used as inputs for the biosynthetic products, as well as in terms of transcribing and regulating enzymes involved in PSM pathways (18). It is unclear, however, whether biosynthetic costs and pleiotropy of biosynthetic enzymes limit phytochemical diversity or lead to evolutionary trade-offs between chemical classes (19–21).

Here we ask how plants generate chemical diversity and what evolutionary processes lead to novel compounds and unique chemical profiles. The evolution of novel chemical profiles is often assumed to be the result of stepwise changes to chemical structures resulting in more derived chemical defenses over evolutionary time (2, 22). Extensive reviews show that most secondary metabolites originate from a small group of precursor compounds derived from primary metabolism, with gene duplication playing a fundamental role (23, 24). Duplicated genes can alter expression levels as well as accumulate mutations leading to neofunctionalization (e.g. changes to substrate specificity) (23, 24). In addition, transcriptional regulation of biosynthetic pathways can lead to novel combinations of existing compounds and the rapid generation of novel chemical profiles (23). While phytochemical diversity is likely generated in part by all of these molecular mechanisms, it is unclear if evolution proceeds through gradual changes that accumulate over evolutionary time, leading to closely related species with similar chemical profiles or if abrupt changes facilitate the rapid divergence of chemical profiles irrespective of shared evolutionary history.

To understand the processes that underlie the evolution of phytochemical diversity we focus on the neotropical tree genus *Inga* (Fabaceae) as a case study for how secondary metabolism evolves within a single lineage. *Inga* is a speciose genus with ∼300 tree species in tropical moist forests throughout the New World. At any given site, it usually constitutes one of the most abundant and speciose genera, with more than 40 coexisting species (25). Multiple lines of evidence have implicated the importance of chemistry in the ecological and evolutionary processes that have shaped the genus (3, 7, 26). Moreover, *Inga*, and other speciose tropical genera such as *Bursera, Psychotria, Piper* and *Protium* are among the most phytochemically diverse plant lineages that have been documented, often having more compounds in a single genus than entire plant communities in temperate ecosystems (27). Thus, *Inga* is an illustrative model for the generation of phytochemical diversity as a whole.

## Materials and Methods

### Study sites and species sampling

We sampled *Inga* between 2005 and 2014 at five lowland tropical rainforest sites (Table S1) spread across the distributional range of the genus. We spent approximately 16 people-months per site collecting data in the field. Specifically, we exhaustively searched each site for all *Inga* species, taking measurements on morphological and defense traits. Species identifications were based on the combination of morphology, phylogenetic reconstruction (28), and in some cases for morphologically difficult to distinguish species we relied on chemocoding (29). Young leaves at approximately 50% full expansion were collected from 5 to 10 spatially separated individuals (with very few exceptions for rare species where we included 3 individuals) in the understory. In general we found the chemical profile of each species to be highly canalized and previous work has shown that 5 individuals is sufficient to capture about 75% of compounds encountered in up to 15 individuals (17). Samples were dried at ambient temperature *in silica* immediately following field collection, and then stored at -20°C.

### Characterization of *Inga* Chemistry

#### a) Soluble secondary metabolites

Metabolites were extracted from dried leaf samples in the Coley/Kursar lab at the University of Utah using a solution of (60:40, v/v) ammonium acetate buffered water, pH 4.8: acetonitrile, producing 2mL of retained supernatant from 100mg (+/-2.5 mg) of sample for chromatographic analysis (Wiggins et al., 2016). Extraction weight (percent dry weight) was measured gravimetrically by subtracting dry marc from the mass of pre-extraction plant material. Small molecules (detector range of 50-2000 Da) from the extraction supernatant were analyzed using ultraperformance liquid chromatography (Waters Acquity I-Class, 2.1 × 150mm BEH C18 and 2.1 × 100 mm BEH Amide columns) and mass spectrometry (Waters Xevo G2 QToF) (UPLC-MS) in negative ionization mode. MS/MS spectra were acquired by running DDA, whereby MS/MS data were collected for all metabolites that ionized above a set threshold (5000 TIC).

#### b) L-Tyrosine

Some *Inga* species invest in the overexpression of the essential amino acid L-tyrosine as an effective chemical defense (30). Tyrosine is insoluble in our extraction buffer, so a different protocol was used to determine the percentage of leaf dry weight. Following (30), extractable nitrogenous metabolites were extracted from a 5 mg subsample of each leaf using 1 mL of aqueous acetic acid (pH 3) for 1 h at 85°C. Fifteen microliters of the supernatant were injected on a 4.6 × 250 mm amino-propyl HPLC column (Microsorb 5u, Varian). Metabolites were chromatographed using a linear gradient (17–23%) of aqueous acetic acid (pH 3.0) in acetonitrile over 25 min. Mass of solutes in each injection were measured using an evaporative light scattering detector (SEDERE S.A., Alfortville, France). Tyrosine concentrations were determined by reference to a four-point standard curve (0.2– 3.0 mg tyrosine/mL, r^2^=0.99) prepared from pure tyrosine.

#### c) Compound separation, annotation, and assignment to species

Following HPLC and UPLC-MS data acquisition, metabolites were quantified and assigned available structural information in all samples using an untargeted metabolomics pipeline developed by our research group. In this pipeline, spectral features are extracted from raw MS data, and related features are grouped into compounds. MS/MS data for each compound are then uploaded to GNPS (31) in order to annotate putative structures and calculate spectral similarity between compounds. These analyses generate 1) a species by compound abundance (MS-1 peak intensity measured by total ion current) matrix and 2) a compound by compound MS/MS spectral cosine similarity matrix, which are then combined into a pairwise species similarity matrix which accounts for both shared compounds between species and the MS/MS structural similarity of unshared compounds (see Endara et al. (17) for details). All code for this pipeline is deposited in a git repository (Forrister & Soule, 2020; https://gitlab.chpc.utah.edu/01327245/evolution_of_inga_chemistry).

#### d) Indices for chemical similarity and phytochemical diversity

To test for phylogenetic signal of the entire chemical profile and quantify divergence between species, we developed a method for quantifying overall chemical similarity between two species (17). This provides a challenge because few compounds are shared between species, making classic distance metrics such as Bray-Curtis uninformative (17, 32). Our method, which is similar to but distinct from one developed by Sedio et al. (32), accounts for the fact that two species may have different compounds that are structurally similar (4, 7, 17). Specifically we leverage MS/MS spectra as a proxy for structural similarity between compounds (31). In this method, total chemical similarity between species is a function of the normalized abundance of shared compounds plus the normalized abundance of unshared compounds weighted by their structural similarity in the molecular network (see (17) for details).

We quantified investment in phytochemical diversity for each focal species using its chemical profile and the MS/MS molecular network to calculate the functional Hill number (33). This diversity measure accounts for both variation in compound abundance and structural similarity in the molecular network. In short, it calculates the effective number of equally abundant and structurally distinct compounds produced by a given species (33). We compared this diversity index with a null model where we assembled compounds into chemical profiles through a bifurcating process from root to tip on the *Inga* phylogenetic tree. The null model represents the chemical profiles randomly drawn from the entire chemical space (see Appendix 1 for detailed explanation of the null model). To make a representative null model we matched the number of compounds produced by a given species and the number of compounds shared between any two closely related species with the values observed in the actual data. We normalized phytochemical diversity values of each species relative to our null model.

### Phylogenetic reconstruction of *Inga*

A phylogenetic tree containing 165 *Inga* accessions, including the taxa sampled at multiple sites, was reconstructed using a newly generated targeted enrichment (HybSeq) dataset of 810 genes. These 810 loci include those presented in Nicholls et al.(28), supplemented with a subset of the loci from work by Koenen et al. (34). DNA library preparation, sequencing and the informatics leading to final sequence alignments follow protocols in Nicholls et al. (28). For the phylogenetic inference, we accounted for the putative effect of incomplete lineage sorting by constraining the maximum likelihood phylogeny with the topology obtained from a coalescent-based method. First, we inferred gene trees for 810 loci using *IQtree* 2 (35). The best substitution model was estimated for each loci using the ModelFinder (36) module implemented in IQtree 2. For each gene tree, we performed 1,000 bootstrap replicates with the ultrafast bootstrap approximation (37). The resulting gene trees were subsequently used as the input for ASTRAL-III to estimate a phylogeny in a summary coalescent framework (38), after contracting branches with bootstrap support <10. We then used the topology obtained with ASTRAL to perform a constrained maximum likelihood tree search in IQtree 2. We performed a partitioned analysis (39) after inferring the best-partition scheme for the 810 genes and the best substitution model for each partition using ModelFinder. Branch support was estimated with ultrafast bootstrap approximation (1,000 replicates). The phylogenetic tree was subsequently time-calibrated using penalized likelihood implemented in the program treePL (40). We used cross-validation to estimate the best value of the smoothing parameter and implemented secondary calibration points on the crown and node ages of *Inga* with an interval of 9.2-11.9 My and 13.4-16.6 My, respectively. Finally, the complete phylogeny was pruned to include only the 98 species for which chemistry data were available.

### Phylogenetic Comparative Methods & Ancestral State reconstruction

For phylogenetic signal of continuous traits we calculated Blomberg’s *K* (41) using function *phylosignal* in the R package *picante* v.1.8.2 (42). *K* is close to zero for traits lacking phylogenetic signal, and higher than 1 when close relatives are more similar than expected under the classic Brownian motion evolutionary model. For the presence and absence of individual compounds we calculated the D-statistic (43) using the *caper* package (44). We took a stochastic character mapping approach for the ancestral state reconstruction of compound presence absence on the *Inga* phylogeny. Specifically, we used the function *make*.*simmap* (45) from R package *phytools v*.*0*.*7-47* (46) to estimate the state of each internal node on the phylogeny using 100 simulated trees.

## Results

We studied *Inga* at five sites across the Amazon basin and in Panama (Table S1), where we extensively surveyed understory saplings, a prolonged and key vulnerable stage in the life cycle of tropical forest trees (26). We identified all species of *Inga* present at each site and characterized the chemical profile of expanding leaves for a total of 97 species as well as one species from its sister genus, *Zygia*. We focused our study on the chemical defenses of expanding leaves, as they receive more than 70% of the lifetime damage of a leaf (47), and their chemical profiles are an important factor for host associations of insect herbivores (3, 4).

We developed methods for characterizing the complete phytochemical profile of expanding leaves using untargeted metabolomics (see methods for details), which allowed us to characterize thousands of individual compounds and determine the similarity of chemical profiles across species. Our untargeted metabolomics pipeline revealed 9,105 unique compounds across 808 samples. *Inga* species invest substantial resources in chemical defense, averaging 194 ± 103 (mean ± s.d.) unique compounds per species, and comprising 37 ± 11% (mean ± s.d.) of the expanding leaf’s dry weight (Fig. S1). We employed a variety of techniques in order to assign individual compounds into classes including NMR structural characterization, MS/MS-based spectral library searches, *in silico* compound annotation, and machine learning prediction. As a result, we were able to classify 42.5% of compounds, a vast improvement from the 2.9% achieved from library matches alone (Fig. 1). Although our extraction and detection methods did not explicitly exclude primary metabolites, the vast majority of annotated compounds were assigned to secondary metabolites, specifically chemical classes that have been classically implicated in plant defense against pathogens and herbivores, including flavonoids and saponins. Moreover, primary metabolites are generally observed in much lower concentrations than secondary metabolites and thus would not be readily detected in our UPLC-MS pipeline. Finally, when these chemical extracts were incorporated at only 0.5–2% DW into artificial diets, they were highly detrimental to larval growth and survival, suggesting that they are toxic (reviewed in 24).

**Figure 1:**
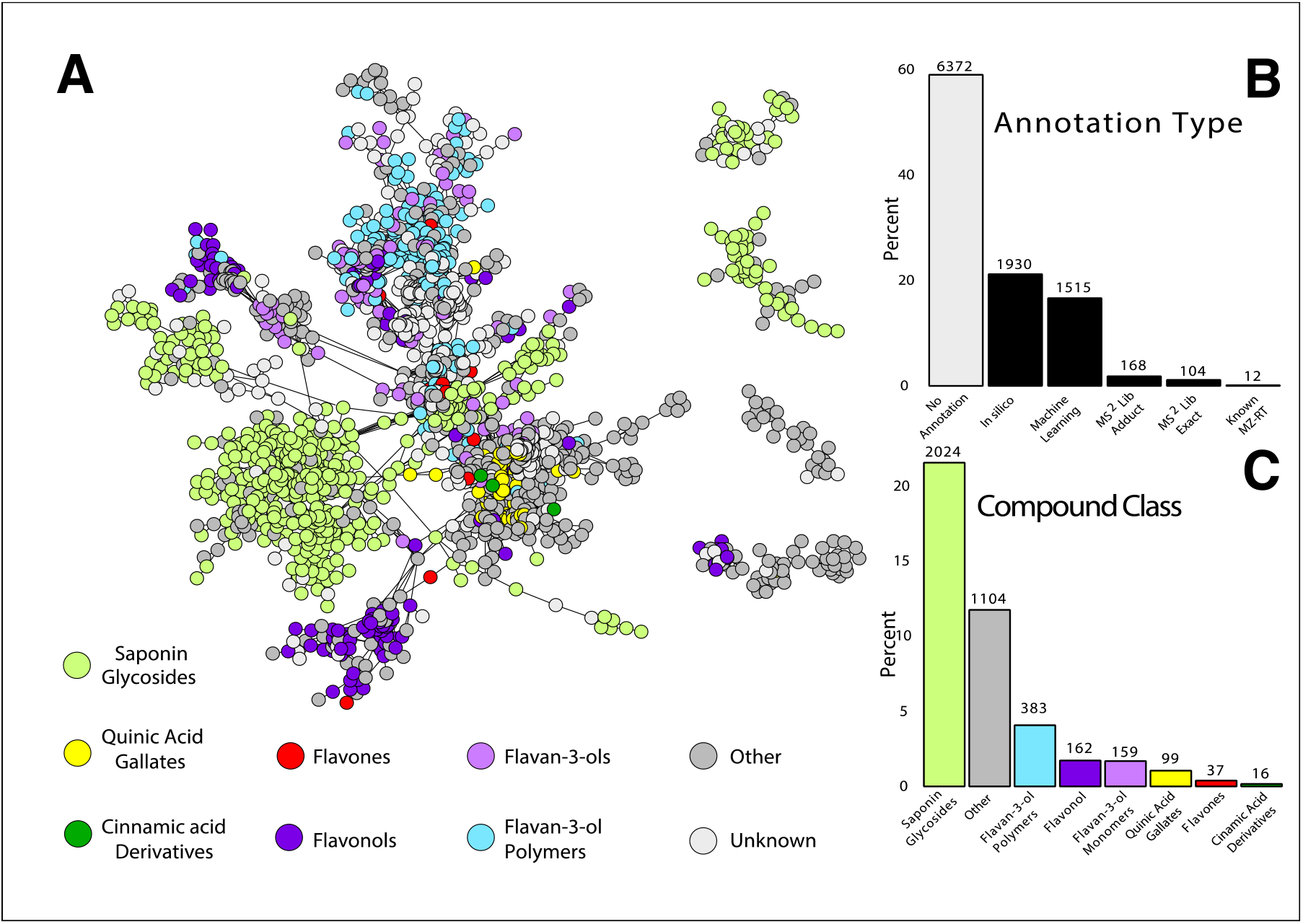
Compound based molecular network: (A) Subset of molecular network (see Fig. S2. for the full network) containing all compounds observed across 98 study species. Nodes represent individual compounds identified in the metabolomics pipeline, and connections between compounds (edges) are based on the MS/MS cosine similarity score from GNPS (https://gnps.ucsd.edu). (B) Percent of compounds that were annotated using different methods -*in silico* fragmentation, machine learning, MS/MS library exact matches and adducts, and comparison to authentic standards on our UPLC-MS system based on mass-charge ratio (*m/z*) and retention time (RT). (C) Percent of compounds with annotations represented by each compound class. For B and C, total number of compounds are reported at top of bars.

We leverage this metabolomic dataset along with comparative phylogenetic methods to explore the means through which phytochemical diversity evolves, addressing the following questions:

### Do species maximize phytochemical diversity, or are they constrained by biosynthetic costs, trade-offs or genetic constraints?

Investment in structurally diverse defensive compounds is adaptive for protection against a broad suite of herbivores (8, 11, 12), yet investment in chemical defense comes at a cost (known as the ‘growth-defense trade off’) (48–50). We ask whether biosynthetic tradeoffs constrain a plant’s ability to invest in structurally unrelated compounds (i.e., the cost of maintaining enzymes in multiple PSM pathways), or whether selection promotes investment in chemical diversity.

To answer this question, we quantified investment in phytochemical diversity using functional Hill numbers and compared these findings to a null model. For more than half of the species (60% of the species), phytochemical diversity was within the range of values expected by our null model. The rest of the species exceeded that range (38% of species; Fig. 2). The absence of species with lower phytochemical diversity than the null model suggests that any biosynthetic or genetic constraints that would limit diversity are overcome by the apparent strong selection by pests and pathogens for phytochemical diversity.

**Figure 2:**
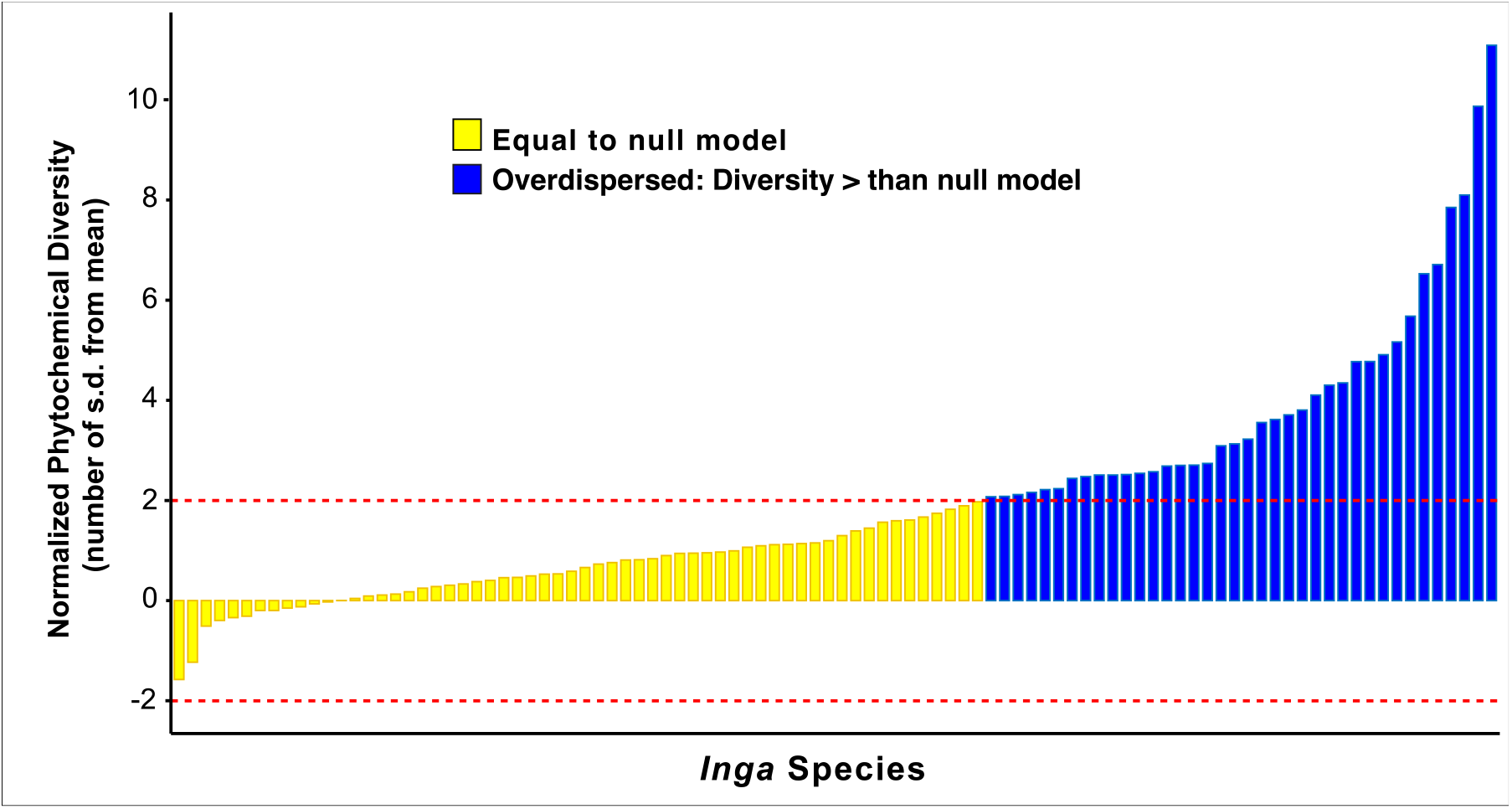
Normalized phytochemical diversity in each *Inga* species: Bars represent individual *Inga* species ordered by increasing phytochemical diversity measured as the functional Hill number. Values represent the number of standard deviations above or below the mean calculated in the null model, with dashed red lines indicating 2 standard deviations above and below the null mean. Values less than zero represent species that are less chemically diverse than a random draw (under-dispersed in the MS/MS network) and values above zero represent species that are more diverse (over-dispersed in the MS/MS network).

### Does the entire chemical profile diverge between closely related species?

The classic ‘escape and radiate’ model predicts that closely related species would have similar defensive profiles (2, 30, 51). However, it has also been posited that diffuse coevolution between plants and their natural enemies would result in divergent adaptation in defense traits (52, 53). Thus, it is advantageous for a species to not only have a diversity of compound classes, but also to be different from other species in their community in order to not share herbivores (7, 12, 16). Here we ask if species’ chemical profiles show phylogenetic signal, or if they have diverged sufficiently to erase the effect of shared evolutionary history. Previous work in *Inga* shows that defense strategy has little phylogenetic signal (3, 16), a pattern that has been documented in other plant lineages (12, 54, 55). However, previous work focused on the evolution of a few specific metabolites (30) or broad compound classes (16). In this study, we leverage metabolomics to greatly expand our exploration of the relationship between evolutionary history and chemical similarity by tracking the relative abundance of over 9,000 individual compounds. Moreover, we use a novel modeling framework to formally test the hypothesis that chemical profiles are evolving under divergent adaptation.

To test for phylogenetic signal of the entire chemical profile and quantify divergence between species, we developed a method for quantifying overall chemical similarity between two species (17). Lastly, we compare these calculations to estimates of chemical similarity we would expect from a null model (described in methods and appendix 1). We found that chemical similarity was highest for intraspecific comparisons, but quickly decreased to the point where two species were as dissimilar as expected under our null model for all interspecific comparisons (Fig. 3; Fig. S3). Within a species, chemical similarity was highest between individuals at a single site (but rapidly decreased between individuals of the same species at different sites (Fig. 3). Population-level chemical divergence occurs despite the fact that there is essentially no limitation on the dispersal of *Inga* species across the Amazon, such that the metacommunity for any site is the entire Amazon basin (17, 56). Between-site differences in abiotic and biotic conditions may drive intraspecific population-level differences in chemical profile, including variation in soil types and precipitation patterns or the complete turnover of herbivore communities.

**Figure 3:**
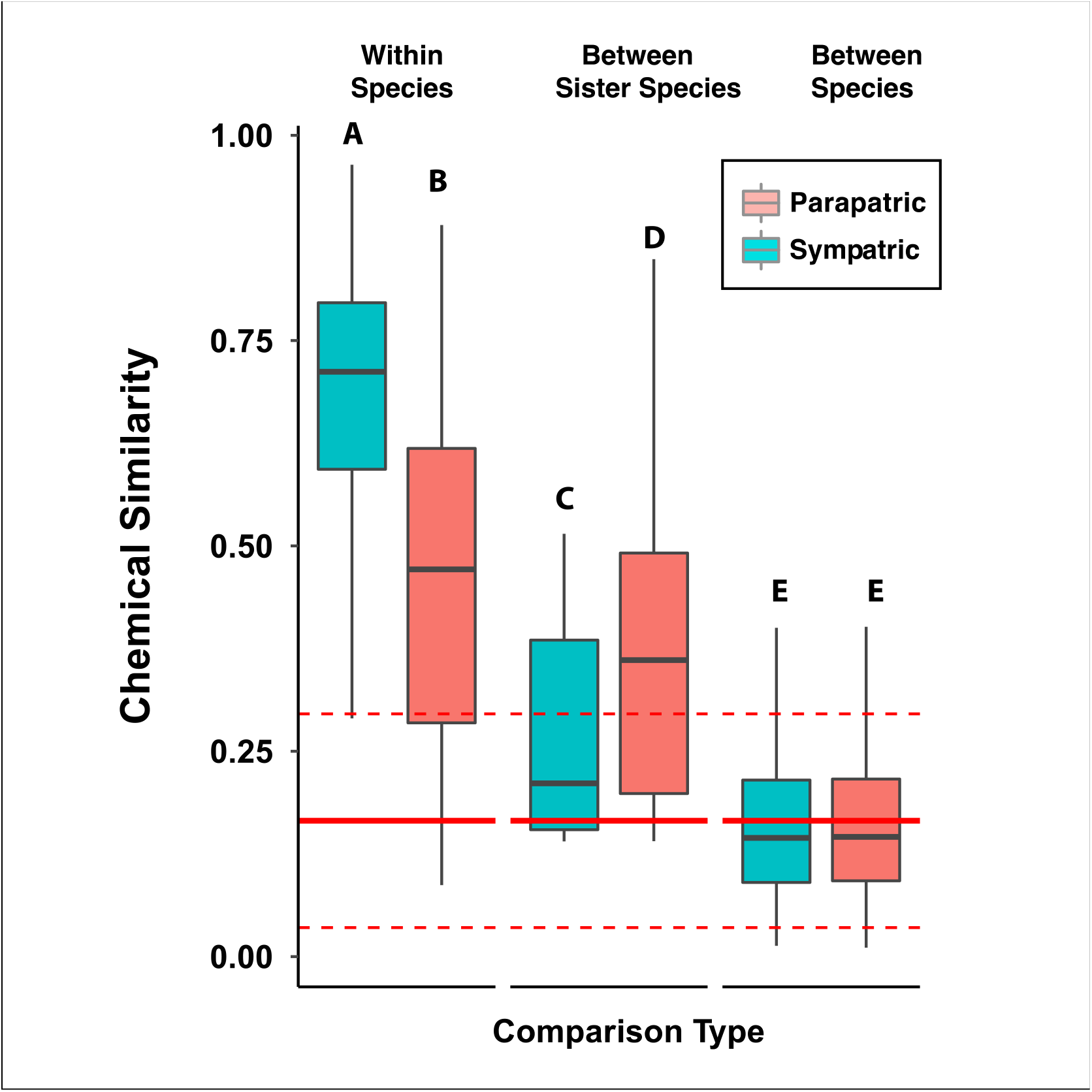
Comparison of entire chemical profiles between *Inga* Species: A) Boxplot comparison of chemical similarity scores for *Inga* within a species, between sister species, and between all other species. Comparisons between and within sites are indicated by red and blue boxes, respectively. Significantly different groups are denoted by A, B, C, D, and E. The solid red line indicates the mean chemical similarity score observed in the null model which simulates the expected chemical similarity between two randomly assembled chemical profiles. The dashed red lines represent 2 standard deviations above and below the null mean.

We also found that interspecific chemical similarity was highly divergent even between sister species and that the majority (83%) of pairwise comparisons between species fell within the range of our null model (Fig. 3, Fig. S3). Sister species at different sites (parapatric) were divergent, presumably because of between-site differences in selection pressures. However, surprisingly, sympatric sister species were more divergent than parapatric sister species. This suggests that that it is adaptive for sister species exposed to the same herbivore community to diverge in defenses, highlighting the role of ecological niche divergence in species coexistence. Interspecific chemical similarity of the entire chemical profile showed no phylogenetic signal (Mantel test: r= -0.03, P= 0.68, Fig.S3), providing further evidence that chemical profiles rapidly diverge and are unconstrained by shared evolutionary history. We posit that antiherbivore defenses, including chemical defense, are one of the first traits to diverge during the speciation process, especially compared with non-defensive traits such as those used for resource acquisition (57).

### Is defensive chemistry evolving under divergent selection?

To formally test the hypothesis that a species chemical profile is evolving under divergent selection, we used recently developed phylogenetic comparative methods to model different modes of trait evolution and select the best fitting model. We compared Brownian motion, Ornstein–Uhlenbeck and a newly developed model for divergent adaptation, which adds a term for interactions between lineages during trait evolution (58). We found strong support for the divergent adaptation model over models that assume all lineages evolve independently of others on a tree (i.e. Divergent vs, Brownian motion and the Ornstein– Uhlenbeck process) (Table S2). Our results show that each species evolves to have a unique chemical profile compared to close relatives, likely driven by divergent selection imposed by insect herbivores and other natural enemies. Unlike a species chemical profile, we found that traits related to the amount of chemical investment (number of compounds, chemical investment and phytochemical diversity; Fig. S1) were best explained by an Ornstein–Uhlenbeck process model, indicating that these traits are evolving towards an optimal trait value (Table S2) rather than diverging.

### Are individual compounds phylogenetically conserved?

We now turn to evaluating the underlying evolutionary processes and biosynthetic mechanisms that facilitate the evolution of diverse and unique chemical profiles. The widely accepted ‘escape and radiate’ model for defense evolution was first suggested a half century ago by the work of Dethier (59), Fraenkel (60), and Ehrlich and Raven (2). In the model, random mutations in structural genes occasionally lead to the production of novel defense compounds, often through the gradual embellishment of core structures into more complex and derived compounds (30, 51). If these compounds have stronger deterrent properties or are effective against different enemies, selection acts to promote the novel genotype. Nonetheless, they should still show a pattern of phylogenetic conservatism (12). To test this prediction, we mapped all individual compounds present in *Inga* onto the phylogeny and estimated their phylogenetic signal. We then used ancestral state reconstruction to estimate the number of times each compound had transitioned on the phylogenetic tree (61).

The majority of compounds are detected in only a few species (median = 4), and roughly half (53%) of compounds showed no phylogenetic signal (Fig. 4A). Although some compounds are clustered in specific clades, many compounds are found dispersed across the phylogeny (Fig. 4B). Based on the ancestral state reconstruction of each compound, we created an index of evolutionary lability, calculated as the number of times a given compound transitioned between present and absent divided by the number of species where a compound is present. Low values for this index indicate strong phylogenetic conservatism, where a compound likely evolved few times and was retained within a given lineage. Values near or above 1 indicate that a compound is evolutionarily labile, having been gained or lost frequently throughout the phylogenetic tree. We found that the majority of compounds (58%; lability >= 1.0) were labile having evolved as many or more times than they were present (Fig. 4C). Although compounds spread throughout the phylogeny could have evolved independently by convergent evolution, the scale of how frequently compounds are apparently gained and lost is more consistent with the up– and down-regulation of key enzymes via transcriptional regulation (23, 61).

**Figure 4.**
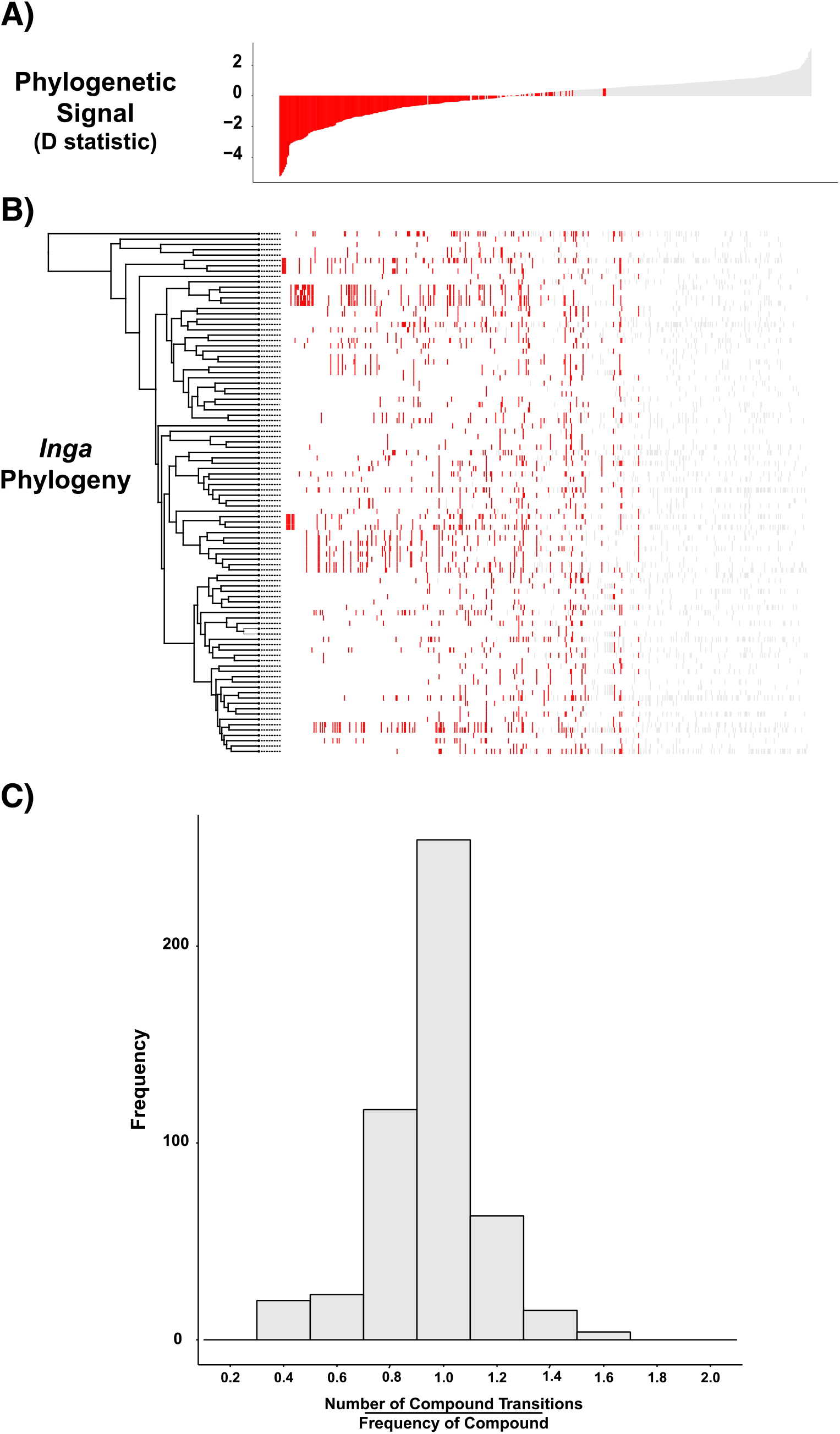
Expression patterns of individual compounds mapped onto the *Inga* phylogeny: (A) Phylogenetic signal of 500 randomly sampled compounds ordered from most to least phylogenetically conserved using the D statistic. For visualization purposes we display 500 randomly chosen compounds. Red bars indicate compounds with significant phylogenetic signal (p <0.05). (B) Heat map demonstrating expression of individual compounds on the phylogeny. Red (significant phylogenetic signal) and grey (non-significant) bars indicate where a compound is present in a given species. (C) Histogram for the compound lability index for all compounds present in > 2 species.

### Is there evidence of metabolic integration or apparent trade-offs between biosynthetic pathways?

Comparative phylogenetic analyses of defense traits have revealed both trade-offs (negative correlations) (19, 26, 50, 62, 63) and positive correlations (63), providing evidence for evolutionary integration and defense syndromes. For example, trade-offs between compound classes that share the same biosynthetic precursor are well supported in the literature (19, 64, 65). Nevertheless, other studies have found little evidence for these trade-offs based on meta-analysis (21) Here we ask whether physiological constraints lead to trade-offs that persist over evolutionary timescales or if each branch of the biosynthetic pathway evolves independently. To test this, we grouped individual compounds into broad chemical classes based on their biosynthetic origins (Fig. S4A; Fig. S5). We tested for phylogenetic signal of each of these classes and then used phylogenetic structural equation modeling (SEM) to determine if chemical classes were correlated with each other (Fig. S4B). We applied this approach because it controls for the phylogenetic non-independence of species as well as the biosynthetic non-independence of predictor variables.

The chemical profiles of *Inga* species are dominated by two classes of compounds that can be broadly categorized as phenolics and saponins. Phenolic chemistry arises from the flavonoid pathway (Fig. S5 contains a summary of *Inga* phenolics). The majority of *Inga* phenolic chemistry is based on flavone and mono/polymeric flavan backbones that are extensively modified. *Inga* saponins are glycosylated triterpenoids that have their origin in the mevalonic acid pathway and as such are biosynthetically distinct from phenolic compounds. We mapped investment in each of these classes onto the phylogeny (Fig. S4A) and then tested for phylogenetic signal of each subclass of these compounds. We found that quinic acid gallates (K= 0.68, p = 0.02), tyrosine and related depsides (K= 0.73, p=0.03) as well as saponin glycosides (K= 1.02, p=0.007), showed significant phylogenetic signal. In contrast, all flavonoid subclasses showed no phylogenetic signal (Fig. S4A). The fact that the flavonoid subclasses are not restricted to a given clade suggests that all species are capable of producing these compounds whereas other compound classes likely evolved once or few times and are restricted to specific clades (30).

Our SEM model revealed several trade-offs between compound classes suggesting that there may be switch points between major branches of the biosynthetic pathway: 1) saponin glycosides were negatively correlated with the left and right branch of the flavonoid pathway, 2) quinic acid gallates were negatively correlated with the right side of the flavonoid pathway and 3) the right branch of the flavonoid pathway was negatively correlated with the left branch (Fig. S4B). Although our model revealed moderate trade-offs across pathways, it should be noted that many species of *Inga* produce compounds from multiple biosynthetically distinct compound classes (Fig. S4B). This is consistent with our findings that *Inga* invest in phytochemical diversity (Fig. 2) and with a meta-analysis that found little evidence for trade-offs in defensive traits (21). The ability for some species to produce compounds from up to five different compound classes coupled with the fact that one class did not completely exclude production of other classes indicates that these trade-offs may not be driven by hard physiological constraints. For example, saponin production was negatively correlated with investment in flavan-3-ols, yet there were nine species that invested in both of these pathways simultaneously. The lack of strong physiological constraints likely facilitates the evolution of novel chemical profiles and divergence between closely related species.

## Discussion

Our findings highlight the exceptionally high levels of phytochemical diversity produced by *Inga* species, despite costs, tradeoffs and constraints of biosynthesis. This result underscores the strong selective pressure by plant natural enemies that would favor investment in a diversity of compounds. Furthermore, in contrast to the conventional model, closely related species have divergent chemical profiles, with most individual compounds and chemical classes showing little phylogenetic signal and exceptionally high lability across the genus. Divergence of biosynthetic pathways would permit sister species to present very different chemical profiles, reducing the probability that they will share natural enemies. The fact that we observed divergent chemical profiles between close relatives in parapatry, is unsurprising given many differences across sites in abiotic and biotic selection pressures (66). However, the fact that sister species in sympatry (where all individuals are exposed to a similar community of pests and abiotic conditions) displayed much higher niche divergence (Fig. 3), is consistent with natural selection to not share pests (7, 67). Thus, the evolutionary fluidity of defensive chemistry may be a major factor allowing long-lived trees to effectively persist in the arms race with insect herbivores.

Increasingly, evidence is supporting the adaptive value of chemical diversity both within and among plant species. But how are novel structures generated and what is the mode of chemical evolution? It is widely accepted that most secondary metabolites originate from a small group of precursor compounds derived from primary metabolism with gene duplication and neofunctionalization driving the modification of basic core structures (22). Classic theory (2), as well as evidence from studies on several plant lineages, predict that stepwise biosynthetic embellishments will gradually produce more complex structures over evolutionary time (30, 51). However, for *Inga*, an alternative mechanism for generating novel chemical structures may be operating. In addition to small embellishments on a compound’s core structure, such as the addition of a methyl or hydroxyl group, we commonly see the addition of larger structures, such as phenolic acids and carbohydrates. The addition of these side groups in a combinatorial manner, referred to as “Lego-chemistry,” has been shown to generate an impressively diverse array of larger structures from a small group of building blocks (68, 69).

Lego-chemistry could be particularly important for the generation of novel structures in the phenolic biosynthetic pathway, which produces the most diverse class of compounds in *Inga* (Fig. S5). *Inga* produces several subclasses of flavonoids that are further modified by the addition of divergent combinations of R-groups to key linkage sites on the basic scaffold molecule (flavonoid aglycones). These R-groups include structures like phenolic acids and carbohydrates which are often derived from pathway precursors and intermediates. For example, (epi)catechin (Fig. S5, comp 27), one of the most common compounds in *Inga*, is modified into at least 4 divergent structures (illustrated in Fig. S6), which upon polymerization leads to the generation of at least a dozen unique polymers (Fig. S5, comp 34).

Although both small chemical embellishment and Lego-chemistry provide a mechanism for plants to generate novel structures, the concomitant evolution of regulatory mechanisms (e.g. at the gene expression level) is required to produce a pattern of phylogenetically dispersed expression of individual compounds (Fig. 4). Thus, we hypothesize that the mode of chemical evolution for *Inga* is the combination of Lego-chemistry as a means to generate novel structures along with changes in regulation of gene expression to generate unique chemical profiles in each species.

Changes in gene expression would permit distantly related species to express the same compound and closely related species to express divergent compounds (61). For example, one sister species could make saponins and its close relative could make phenolics, presenting very different detoxification challenges for herbivores. This model of chemical evolution would imply that species maintain a complete set of biosynthetic enzymes within their genome that are up-or down-regulated in different species and that “unused” genes would have to remain functional over evolutionary timescales. Preliminary results from two *Inga* genomes indicate that the core biosynthetic genes involved in flavonoid and saponin biosynthesis are in fact present in all species even when they do not produce these compound classes (pers. comm. C.A. Kidner, 2021). The maintenance of these supposedly unused enzymes may be required by deep homology and pleiotropy for core biosynthetic enzymes (23, 24). We offer several possibilities for how viable genes are maintained. First, many compounds, including pathway intermediates, do not accumulate to physiologically significant levels. However, because they are essential for the synthesis of downstream compounds, the enzymes responsible for them must be transcribed and maintained. This is the case for the phenylpropanoid compounds that link the shikimic acid pathway with the flavonoid pathway (Fig. S5). Second, it is possible that many compounds that are absent in leaves could be present in other tissues (70, 71). Finally, because there are many more secondary metabolites than enzymes that produce them, it has been argued that a core set of enzymes with low substrate specificity is capable of producing a broad set of chemical structures (20, 72). This concept has proven to be important for generating novel structures via Lego-chemistry (20, 72, 73). If this is the case, it is possible that these core enzymes are conserved across all *Inga* species and spatial and temporal pathway regulation alters which products accumulate.

Our goal in laying out the above hypothesized mechanism for the generation of novel compounds and chemical profiles is to integrate the patterns we observed in our study of *Inga* metabolomes, with the growing body of literature on the underlying genetic and biochemical mechanisms for the evolution of plant secondary metabolism (20, 50, 72, 73). We look forward to this model being tested and updated by future studies using multiomics approaches (50).

## Acknowledgments

This work was greatly facilitated by research permits from the governments of Panama, Peru, Brazil and Ecuador and French Guiana. Thanks to Zachary Benavidez, Carine Emer, Julio Grandez, Wilder Hidalgo, Emily Kearny, Mayra Ninazunta, Wilmer Rosendo, Joe Sixto Saldaña, Yamara Serrano, Georgia Sinimbu, Allison Thompson, and Marjorie Weber for their valuable field assistance. This work was supported by the following grants: NSF GRFP to D.L.F.; Nouragues Travel Grants Program, CNRS, France, and the National Science Foundation (DEB-0640630 and DIMENSIONS of Biodiversity DEB-1135733) to P.D.C and the Secretaría Nacional de Educación Superior, Ciencia, Tecnología e Innovación del Ecuador (SENESCYT) to M-J.E.

## Author Contributions

T.A.K., M-J.E., D.L.F., and P.D.C designed and conducted the research. D.L.F. designed and performed the data analysis. T.A.K, D.L.F., A.J.S., G.C.Y., and A.G.M, contributed to the metabolomic analysis. T.A.K. and J.L. provided initial characterization of *Inga* chemistry via NMR. J.A.N., R.T.P., K.G.D., C.A.K., and O.L. contributed the phylogeny of *Inga*. D.L.F., M-J.E., A.J.S. and P.D.C. wrote the manuscript. All authors provided feedback and edited the manuscript.

## Data availability

Chemical data and scripts to estimate chemical similarity are deposited in a git repository (Forrister & Soule, 2020;

https://gitlab.chpc.utah.edu/01327245/evolution_of_inga_chemistry). All scripts for downstream data analysis and figure generation can be found at (Forrister 2021; https://github.com/dlforrister/Evolution_Of_Inga_Chemistry.git)

