## Supplemental Material for "Diversity and Divergence: Evolution of defense chemistry in the tropical tree genus *Inga*"

New Phytologist

***New Phytologist* Supporting Information**

The following Supporting Information is available for this article:

**Fig. S1** Defense investment traits mapped on to the *Inga* phylogeny

**Fig. S2** Expression of major defensive compound classes mapped on the *Inga* phylogeny and SEM based correlation among them.

**Fig. S3** Compound based molecular network containing all compounds observed in 98 study species

**Fig. S4** Correlation between chemical similarity and phylogenetic distance (My) for all interspecific comparisons

**Fig. S5** Biosynthetic context of phenolic compounds in *Inga*

**Fig. S6** Illustration of Lego-chemistry concept

**Table S1**  Site and Sampling information for all 98 study species

**Table S2**  Maximum-likelihood estimates for different evolutionary models of trait evolution

**Methods S1** Code and description of null model for phytochemical diversity and chemical similarity

| Site | Country | Latitude | Longitude | Annual Rainfall (mm) | *Inga* Species (n) |
|---|---|---|---|---|---|
| Barro Colorado Island | Panama | 9°S | 80°W | 2623 | 14 |
| Nouragues | French Guiana | 4°N | 53°W | 3000 | 46 |
| Tiputini (Yasuni National Park) | Ecuador | 0°N | 75°W | 3200 | 41 |
| Los Amigos (Madre de Dios) | Peru | 13°S | 70°W | 2648 | 39 |
| Manaus | Brazil | 2°S | 60°W | 2100 | 29 |

**Table S1** Site and Sampling information for all 98 study species

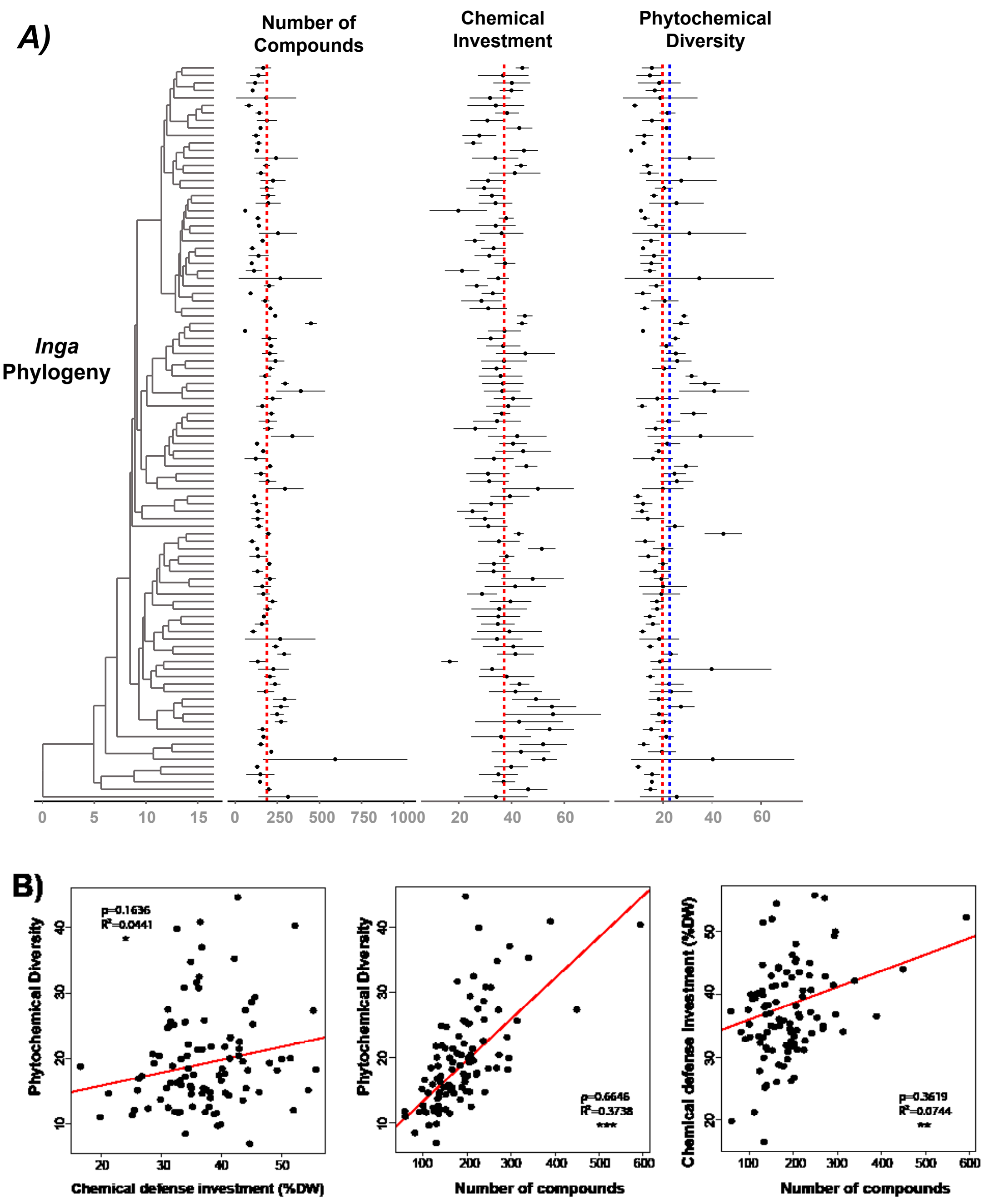

**Fig. S1 A)** Defense investment traits mapped on to the *Inga* phylogeny. Number of unique compounds per species, percent of leaf dry weight invested in secondary metabolism per species, and the phytochemical diversity (measured as functional Hill numbers, q = 2) of each species profile are represented by points. Horizontal bars indicate one standard deviation. Dotted red lines represent mean trait values across all species and the blue line represents the mean value for phytochemical diversity estimated in the null model. **B)** Defense investment trait correlations: (1) Phytochemical diversity vs. percent of leaf dry weight invested in secondary metabolism, (2) phytochemical diversity vs. number of compounds, and (3) percent of leaf dry weight invested in secondary metabolism vs. number of compounds. Points represent individual *Inga* species; red lines represent the phylogenetic linear model estimate of best fit (package: phylolm[1]). Pearson's correlation (ρ), and R-squared are reported, and significance of model fit is represented by asterisks ($p < 0.05$ = *; $p < 0.01$ = **; $p < 0.001$ = ***)

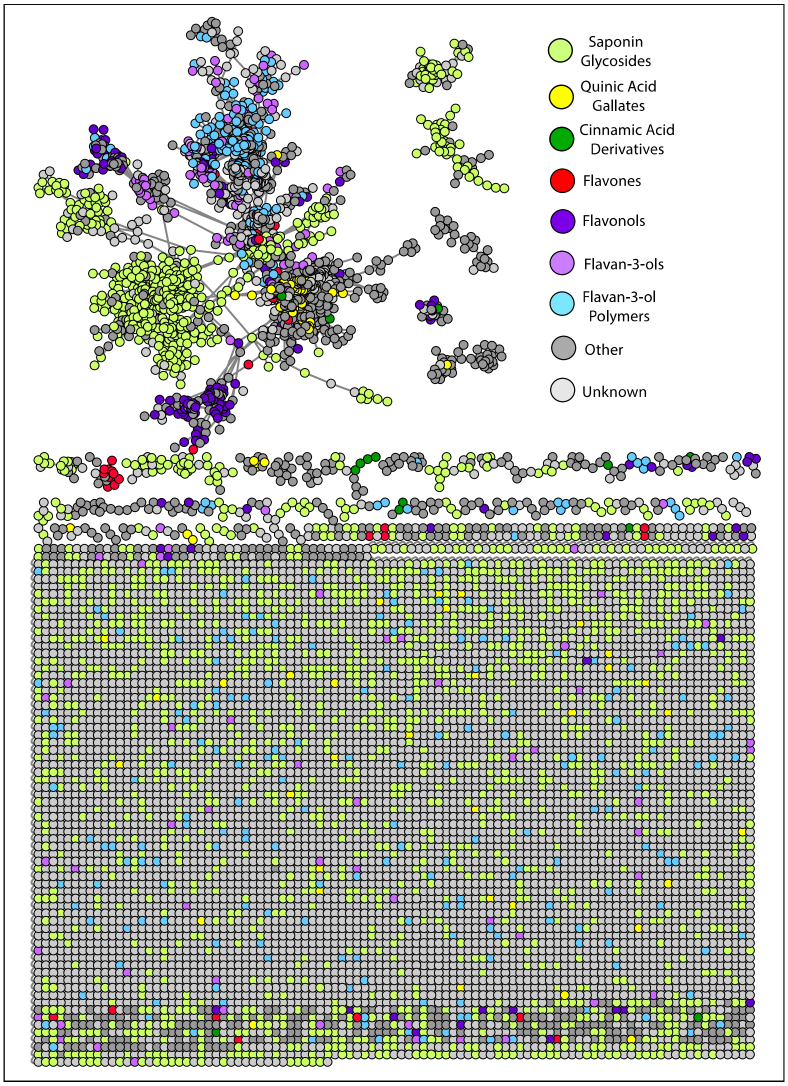

**Fig. S2** Compound based molecular network containing all compounds observed in 98 study species. Nodes represent individual compounds identified in the metabolomics pipeline, and connections between compounds (edges) are based on MS/MS cosine similarity score from GNPS (https://gnps.ucsd.edu). Node color represents compound annotations into major compound classes. Unconnected nodes at the bottom of the network are spectrally unique compounds, that did not match with any other compound in the network, a common feature of ms/ms based metabolomics studies.

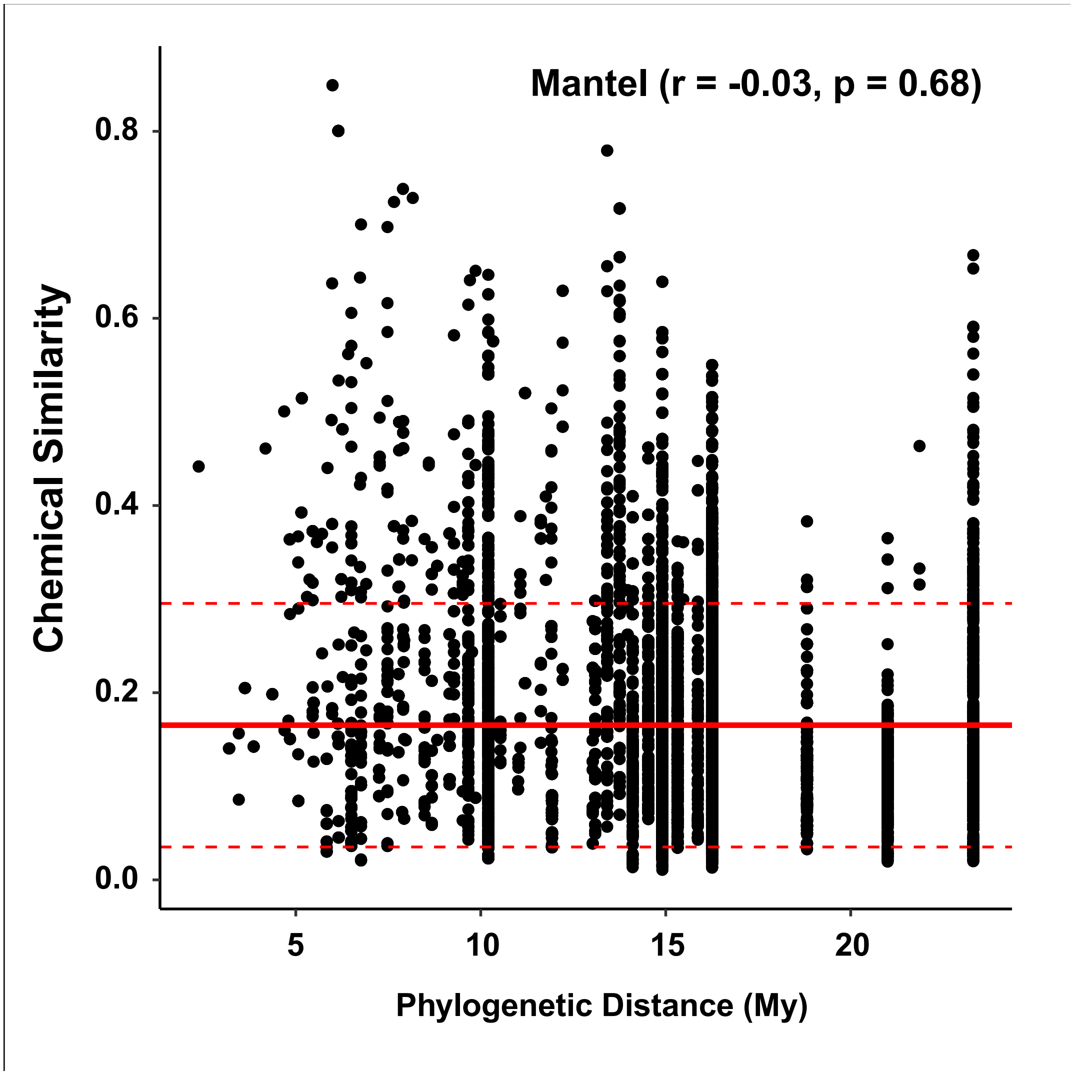

**Fig. S3** Correlation between chemical similarity and phylogenetic distance (My) for all interspecific comparisons. The solid red line represents the mean chemical similarity score observed in the null model which simulates the expected chemical similarity between two randomly assembled chemical profiles. The dashed red lines represent 2 standard deviations above and below the null mean.

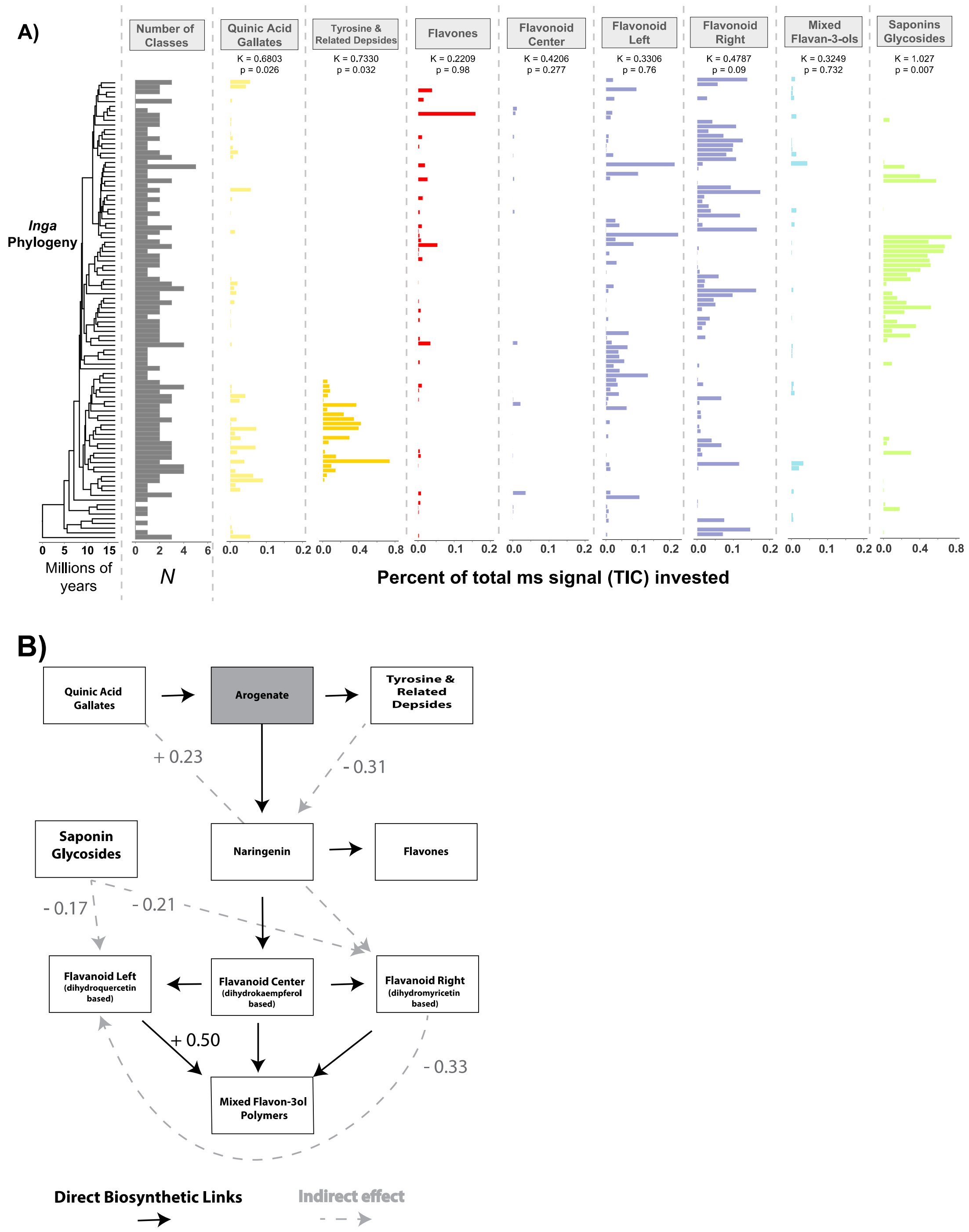

**Fig. S4** (A) Expression of defensive compound classes mapped on the *Inga* phylogeny: Total number of compound classes expressed per species, followed by expression per species of distinct classes including quinic acid gallates, tyrosine and related depsides, flavones, flavonoids, flavan-3-ols, and saponins. Expression of individual compound classes is measured as a percentage of the total MS-level-1 ion current (TIC; metric of abundance) constituted by each class. Phylogenetic signal of each compound class and its significance are represented by Blomberg's K and corresponding p-values. (B) Structural Equation Model (SEM) showing correlation between investment in major defense compound classes produced by *Inga*. Significant ($p < 0.05$) relationships between compound classes indicated by a correlation value listed next to arrows. Solid black arrows represent direct biosynthetic links, regardless of significance of the correlation in the SEM. Dashed grey arrows represent significant indirect relationships between compound classes.

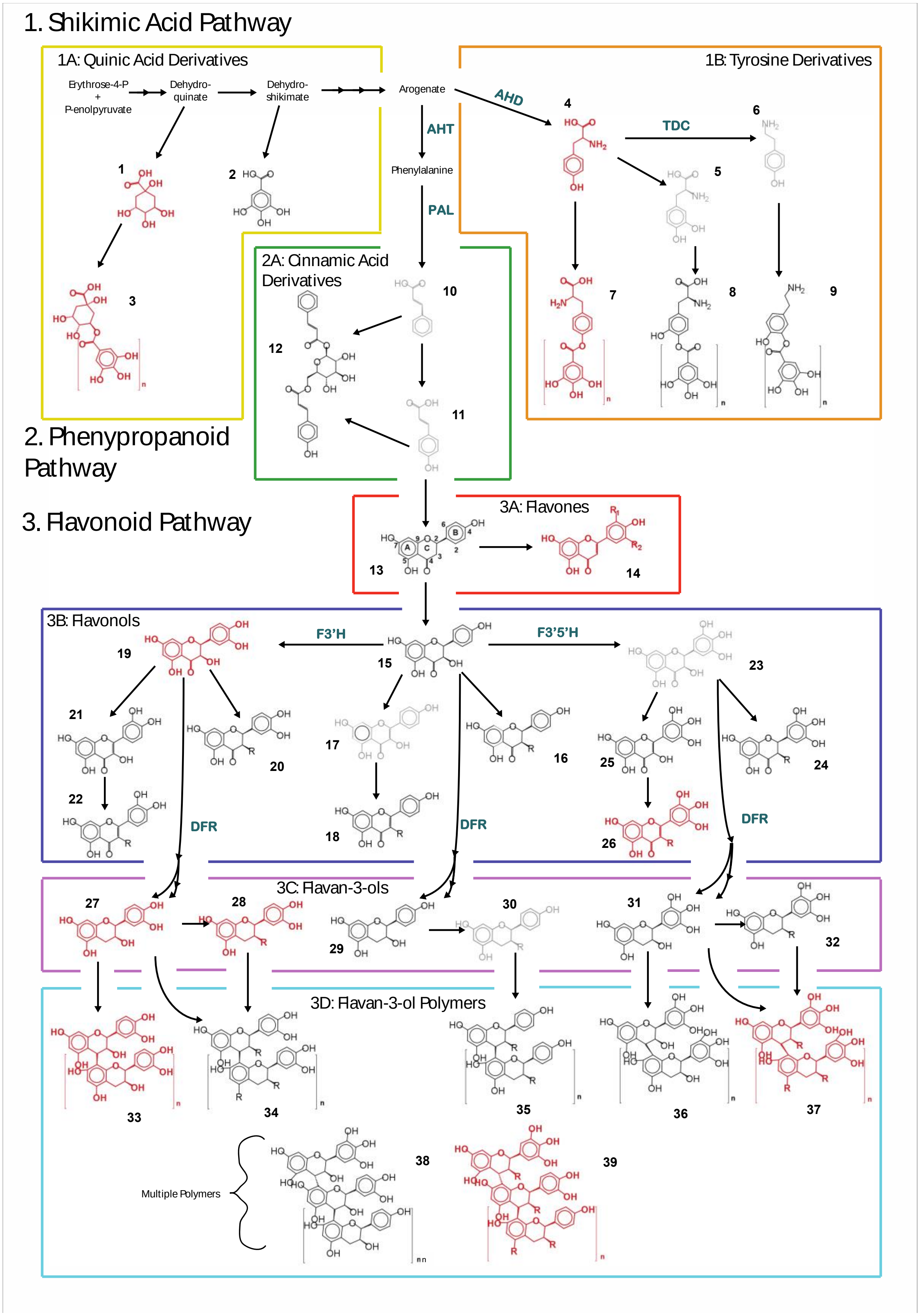
1. Shikimic Acid Pathway
1A: Quinic Acid Derivatives
Erythrose-4-P + P-enolpyruvate
Dehydro-quinate
Dehydro-shikimate
Arogenate
AHD
AHT
Phenylalanine
PAL
1B: Tyrosine Derivatives
TDC
2A: Cinnamic Acid Derivatives
2. Phenypropanoid Pathway
3. Flavonoid Pathway
3A: Flavones
3B: Flavonols
F3'H
F3'5'H
DFR
3C: Flavan-3-ols
3D: Flavan-3-ol Polymers
Multiple Polymers

**Fig. S5** Biosynthetic context of phenolic compounds in *Inga*: (A) Structures and substructures of compounds observed in a survey of 98 focal species and their positions in the biosynthetic pathways that produce them. Compounds that accumulate to significant levels are red and bold; low abundance are black and non-accumulating intermediates are light grey. Wavy bonds indicate variable stereochemistry. Compound names for each compound are listed in Table S3. Marvin was used for drawing, displaying and characterizing chemical structures, substructures and reactions, Marvin 20.20.0, ChemAxon (https://www.chemaxon.com)

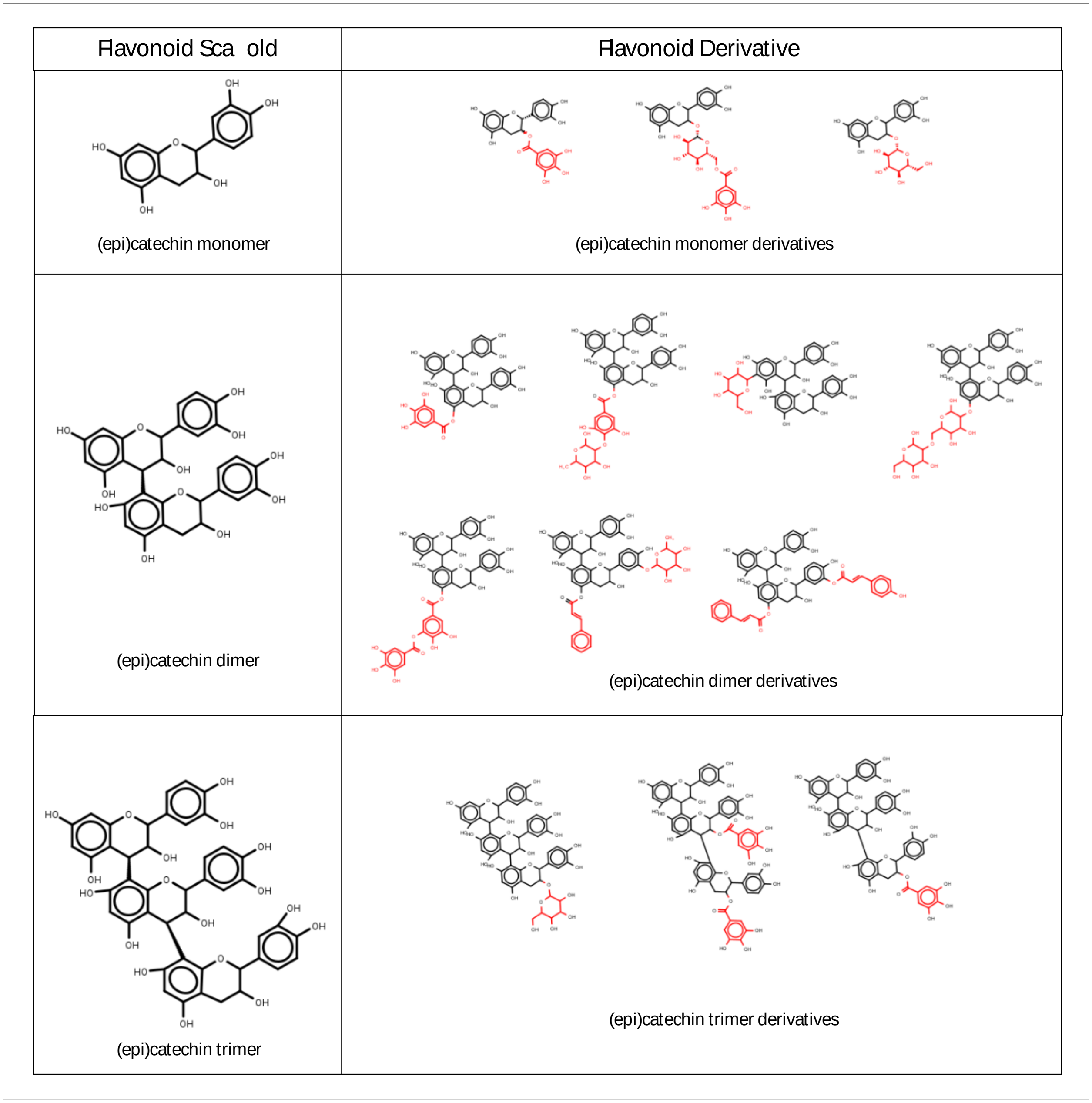

**Fig. S6** Illustration of Lego-chemistry concept based on annotation of monomeric and polymeric Flavan-3-ol compounds observed in *Inga* based on NMR structure elucidation and MS/MS annotation. Red substructures represent commonly observed R-Groups, which are added in a combinatorial manner to generate a variety of compounds.

| Trait | Phyl. Signal | Evol. Model | MLE | Δ AIC | Akaike Weight | P | Interpretation |
|---|---|---|---|---|---|---|---|
| Chemical Profile (Chemotype) | No Phyl. Signal Mantel R = -0.03, p = 0.68 | DA | $\sigma^2$ =14.01, α = 0.24 psi = 0.83 | 0 | 1.0 | *** | Trait lacks phylogenetic signal and is evolving by divergent adapation. |
| | | OU | $\sigma^2$ =0.13, α = 0.13 | 17823 | 0 | *** | |
| | | BM | $\sigma^2$ =0.02 | 17989 | 0 | | |
| Number of Compounds | Significant Phyl. Signal K= 0.56, P= 0.04 | BM | $\sigma^2$ =352.9 | 0 | 0.51 | | Trait shows moderate phylogenetic signal and is evolving under Brownian motion |
| | | OU | $\sigma^2$ =372.43, α = 0.004 | 0.71 | 0.36 | ns | |
| | | DA | $\sigma^2$ = 372.43, α = 0.004, psi = 3.054 | 2.72 | 0.13 | ns | |
| Chemical Investment (% Dry Weight) | Marg. Signif. Phyl Signal K= 0.51, P= 0.06 | OU | $\sigma^2$ =5.02, α = 0.03 | 0 | 0.73 | *** | Trait shows moderate phylogenetic signal and is evolving towards an optimal value. |
| | | DA | $\sigma^2$ =5.02, α = 0.03, psi = 0.00001 | 2 | 0.26 | ns | |
| | | BM | $\sigma^2$ = 3.19 | 66 | 0 | | |
| Phytochemical Diversity | No Phyl. Signal K= 0.36, P= 0.58 | OU | $\sigma^2$ =16.1, α = 0.12 | 0 | 0.72 | *** | Trait lacks phylogenetic signal and is evolving to-ward an optimal value. |
| | | DA | $\sigma^2$ =16.21, α = 0.12 psi = 1.33 | 1.9 | 0.27 | ns | |
| | | BM | $\sigma^2$ =4.34 | 372 | 0 | | |

**Table S2** Maximum-likelihood estimates for different evolutionary models of trait evolution. For each trait we fit three models of trait evolution: A random walk model characterized by Brownian Motion (BM), The Ornstein-Uhlenbeck (OU) model where a trait evolves under BM with a constraining central tendency, and a divergent adaptation (DA) model where trait values the OU model but different lineages interact such that lineage's mean values diverge. We selected the best model based on AIC; significance of model parameters was evaluated by likelihood ratio (LR) tests to determine if a more complex model was significant. Significance indicated by asterisks ($p < 0.05$ = *; $p < 0.01$ = **; $p < 0.001$ = ***).

### Null model for phytochemical diversity and chemical similarity:

Dale Forrister

01/15/2020

Analogous to studies on community assembly, we built a null model in order to put our measure of phytochemical diversity and chemical similarity in perspective. In our model we assembled compounds into chemical profiles through a bifurcating process from root to tip on the Inga phylogenetic tree. We chose this null model because it fixed the number of compounds produced by each species as well as the number of compounds shared between closely related species, while generating chemical profiles randomly drawn from the entire chemical space. Below, we provide the r code for this model as well as key graphs that describe our dataset. We also outline key assumptions assumptions in the model.

##### 1.0 Overview of input data:

##### 1.1 The Inga Phylogeny

This is a species level phylogeny meaning all accessions collected at multiple sites have been combined into a single tip for all sites.

We have 98 species in the phylogeny and it has to be rooted with the Zygia outgroup

```
inga.tree_rooted <- read.tree(here("data",

"Inga_Astralconstrained_datedTreePL_sptree_FINAL_Match_Utl.tre")) plot(inga.tree_rooted,

no.margin = T, edge.width = 2, cex = 0.5, align.tip.label = T)
```

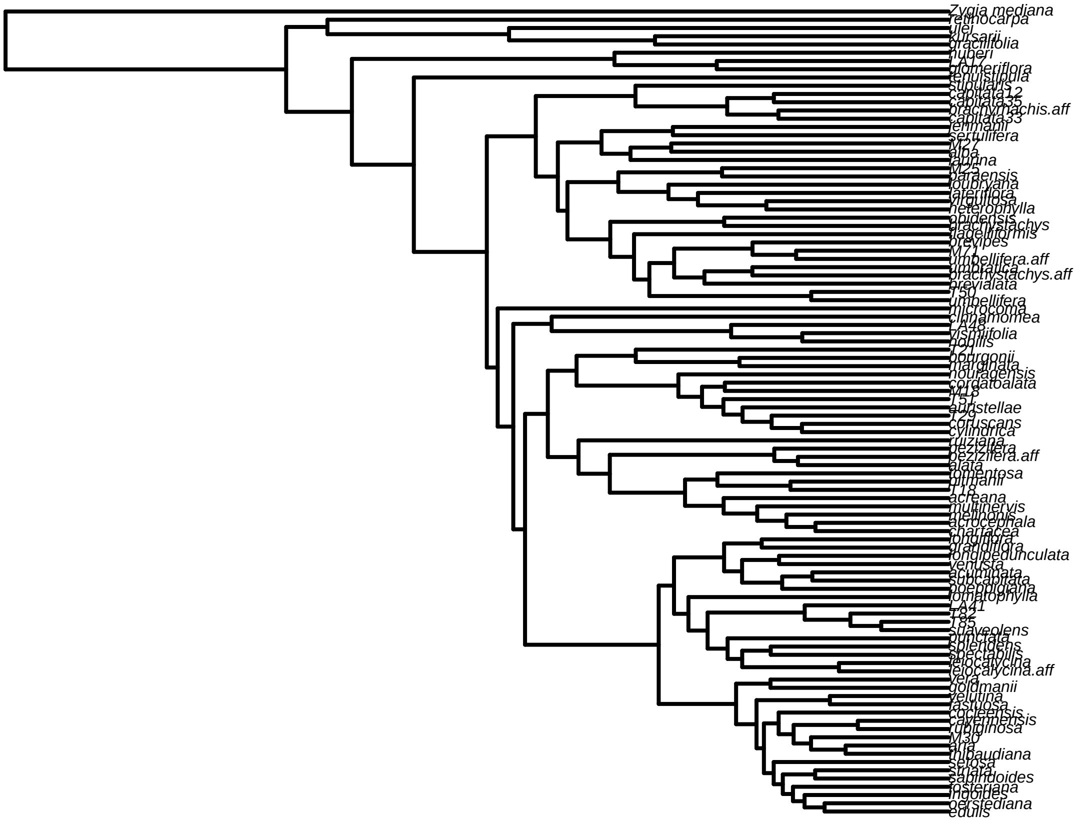

##### 1.2 Inga Chemistry:

Next, we load in chemistry dataset. These include:

1) sampsByCompounds = matrix contain expression of each compound in each sample.
2) pairwise.comps.all = pairwise matrix with cosine similarities between compounds. This is produced by GNPS.
3) pairwise.comps.all_dist = Same thing but converted to distance matrix (1-similarity)
4) phy_code is a key for linking sample names to the phylogeny tip names.

##### 2.0 "Evolving" random samples on a phylogenetic tree:

In our model we assembled compounds into chemical profiles through a bifurcating process from root to tip on the Inga phylogenetic tree. Specifically, we seeded the root node with a chemical profile by drawing at random n compounds from the entire chemical space. We then generated all decedents of this node by: 1) inheriting/passing on a certain percent of compounds in the original profile 2) drawing new compounds, at random such that the new node had n compounds = n compounds observed or estimated in the original data.

```
tree <- read.tree(text = "((Species_1,Species_2),Species_3);")
plotTree(tree, offset = 1) | tiplabels() | nodelabels()
```

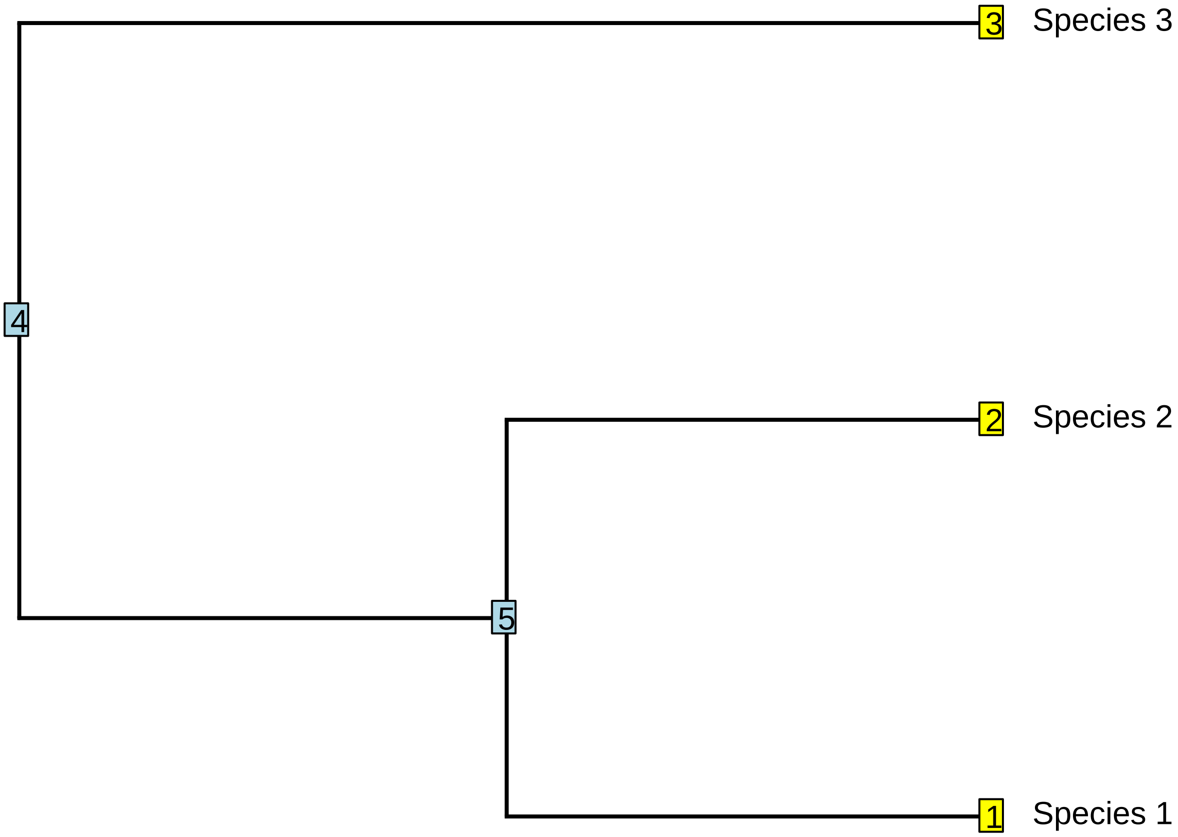

```
## logical(0)
```

For example, in the above simple phylogenetic tree nodes are labeled as blue squares and tips are labeled with yellow squares. We first seed Node 4 with a randomly sampled chemical profile. We then generate Species 3, and node 5. For every node which has decedents, this process is repeated until all tips have been generated. Thus, we then take the generated profile from Node 5 and we modify it to produce Species 2 and Species 1 via the same process.

##### 2.1 Walking through the phylogenetic tree and determining decendents from curent node:

We started by making table of all parents and decedents on the tree. This allows us to walk through the tree generating decedents in the correct order.

```
tree <- inga.tree_rooted
total_nodes <- length(tree$tip.label) + Nnode(tree)
cur_node <- 1 + length(tree$tip.label)

child <- Descendants(tree, cur_node, "children")
while_loop_DF <- data.frame(parent = cur_node, child = child)
nodes_to_do <- data.frame(parent = cur_node, child = child)

while (nrow(while_loop_DF) > 0) {
    node = while_loop_DF$child[1]
    parent = paste("Node_", while_loop_DF$parent[1], sep = "")

    while_loop_DF <- while_loop_DF[-1, ]
    next_nodes <- Descendants(tree, node, "children")
```

```r
    if (length(next_nodes) > 0) {
        while_loop_DF <- rbind(while_loop_DF, data.frame(parent = node, child = next_nodes))
        nodes_to_do <- rbind(nodes_to_do, data.frame(parent = node, child = next_nodes))
    }
}

nodes_to_do$parent <- paste("Node_", nodes_to_do$parent, sep = "")
nodes_to_do$child_name <- NA
nodes_to_do$child_name[nodes_to_do$child > 99] <- paste("Node_", nodes_to_do
$child[nodes_to_do$child >
    99], sep = "")
nodes_to_do$child_name[nodes_to_do$child < 99] <- tree$tip.label[nodes_to_do
$child[nodes_to_do$child <
    99]]

head(nodes_to_do)
```

```
##    parent child    child_name
## 1  Node_99   100      Node_100
## 2  Node_99    98 Zygia_mediana
## 3 Node_100   101      Node_101
## 4 Node_100   193      Node_193
## 5 Node_101   102      Node_102
## 6 Node_101   191      Node_191
```

##### 2.2 Seeding the the root node:

first generate root node by taking a random sample based on the independentswap method using the picante package. Note, to get a sense of variability and stochasticity in our null model, We randomized the entire matrix and then used each row of the matrix as the seed in 98 independent iterations (i)

starting node, is a vector containing the abundance of each compound.

```r
i = 1
sampsByCompounds_rand <- randomizeMatrix(sampsByCompounds, null.model = "independentswap",
    iterations = 10^6)

starting_node <- sampsByCompounds_rand[i, ]

starting_node[1:25]
```

```
##      2      3      5      6      7      8      9     11     12     13     14
##      0      0      0      0      0      0      0      0      0 978045 488904
##     15     16     17     18     19     21     22     23     24     25     26
##      0      0      0      0      0      0      0      0      0      0      0
##     27     28     29
##      0      0      0
```

##### 2.3 inheriting compounds:

The first step in the process of generating a decedent nodes or tips is to inherent some portion of the compounds currently produced in the parent node, defined by the parameters n_comps_shared. See sections 3.1 for details on how this was determined.

To do this we get a list of all compounds present in a parent node and randomly sample ncomps_to_inherit. We then modify the abundance of these compounds by sampling from the actual abundances of that compound found in all samples.

```r
n = 1
all_comps <- colnames(sampsByCompounds)

starting_node <- sampsByCompounds_rand[i, ]
sampsByCompounds_Evolved <- data.frame()
sampsByCompounds_Evolved <- rbind(sampsByCompounds_Evolved, starting_node)

sampsByCompounds_Evolved$node_label <- "Node_99"

sampsByCompounds_Evolved <- sampsByCompounds_Evolved[, c(ncol(sampsByCompounds_Evolved),
    2:ncol(sampsByCompounds_Evolved) - 1)]
names(sampsByCompounds_Evolved) <- c("node_label", names(sampsByCompounds))

sampsByCompounds_Evolved[, 1:10]
```

```
##   node_label 2 3 5 6 7 8 9 11 12
## 1    Node_99 0 0 0 0 0 0 0  0  0
```

```r
child = as.character(nodes_to_do$child_name[n])
parent = as.character(nodes_to_do$parent_names[n])
```

Two functions. . .

```r
inherit_comps <- function(parent, child, starting_node) {
    # if parent = starting node else get parent from evolved comps dataframe
    # (sampsByCompounds_Evolved)
    if (parent == "Node_99") {
        parent_comps <- starting_node
    } else {
        parent_comps <- sampsByCompounds_Evolved[sampsByCompounds_Evolved$node_label ==
            parent, -1]
    }

    # number of compounds in parent node
    parent_comp_ncomps <- sum(parent_comps > 0)

    # determine how many compounds to inherit if child is an internal node we sample
    # from the distribution we find in the actual data child is a tip, we use the
    # actual percent from that node in the transitions data
    if (grepl("Node", child)) {
        ncomps_to_inherit <- round(as.numeric(round(anc_state_ncomps[child])) *
        sample(transitions$per_shared_
            1))
        while (ncomps_to_inherit > length(which(parent_comps > 0))) {

            ncomps_to_inherit <- round(as.numeric(round(anc_state_ncomps[child])) *

                sample(transitions$per_shared_child, 1))
        }
    } else {
        ncomps_to_inherit <- transitions$ncomps_shared[transitions$child == child]
    }

    # sample ncomps_to_inherit from list of compounds present in the parent.
    comps_to_inherit <- sample(names(parent_comps)[which(parent_comps > 0)],
    ncomps_to_inherit, replace = F)
```

```
    # adjust the abundance of each of the compounds present.
    orig_abund_vect <- as.numeric(sampsByCompounds_Evolved[which(sampsByCompounds_Evolved$node_label ==
        parent), -1])
    evolved_abund_vect <- orig_abund_vect

    evolved_abund_vect[which(!all_comps %in% comps_to_inherit)] <- 0

    # for comps that are being inherited sample new abundances based on the
    # abundances in the real data.
    for (comp in 1:length(comps_to_inherit)) {
        species_abundances <- sampsByCompounds[, comps_to_inherit[comp]]
        pot_abundance <- species_abundances[which(species_abundances > 0)]
        replacement_abundance <- pot_abundance[sample(1:length(pot_abundance), 1)]
        evolved_abund_vect[which(all_comps == comps_to_inherit[comp])] <- replacement_abundance
    }
    return(evolved_abund_vect)
}
```

##### 2.4 sampling new compounds

Next, we randomly draw from the entire space to add compounds such that the decedent node contains a total of n compounds. Note the probability of drawing each compound is determined by its frequency in the actual data. See sections 3.2 for details on how this parameter was determined.

```
sample_new_comps <- function(parent, child, evolved_abund_vect) {
    # step 1 sample new comps that aren't already present in order to bring ncomps
    # for that sample up to what it is in the actual data.
    if (grepl("Node", child)) {
        ncomps_to_add <- as.numeric(round(anc_state_ncomps[child])) - ncomps_to_inherit
    } else {
        ncomps_to_add <- transitions$ncomps_new[transitions$child == child]
    }

   new_comps <-
   sample(all_comps[which(sampsByCompounds_Evolved[which(sampsByCompounds_Evolved
   $node_label
        parent), -1] == 0)], prob =
        comp_frequency[which(sampsByCompounds_Evolved[which(sampsByCompounds_Evolved
        parent), -1] == 0)], ncomps_to_add, replace = F)

    # step 2 alter the abundance of the inherited compounds
    for (comp in 1:ncomps_to_add) {
        species_abundances <- sampsByCompounds[, new_comps[comp]]
        pot_abundance <- species_abundances[which(species_abundances > 0)]
        replacement_abundance <- pot_abundance[sample(1:length(pot_abundance), 1)]
        evolved_abund_vect[which(all_comps == new_comps[comp])] <- replacement_abundance
    }

    return(evolved_abund_vect)
}
```

Finally, we then add the new profile to our master list of sampsByCompounds_Evolved.

sampsByCompounds_Evolved <- rbind(sampsByCompounds_Evolved,c(child,evolved_abund_vect))

We then precede to the next row in nodes_to_do, repeating the above two steps until all rows of nodes_to_do have been completed.

##### 3.0 Determining the key pararameters n_compounds_shared and n_compounds total

Our goal is to generate a null model that is directly comparable to the actual data for Inga chemistry. This as much as possible we derive all parameters from the actual values observed in the data.

In our null model we set out to control two aspects of the underlying data 1) fix the number of compounds produced by a given species. 2) Fix the number of compounds shared between two closely related species.

Controlling for these two variables is particularly important when building a null model for the expected chemical similarity between two species. We could have used a more simple null model, in which null chemical profiles are assembled by drawing the mean number of compounds at random from the network. However, this null model would ignore the underlying phylogenetic structure of the data. It is plaussible that evolutionary history plays a role in determine the number of compounds a given compound produces as well as the number of compounds shared between two spaces. By holding these two variables fixed in our null model we hoped to observe the effect of which compounds are in each species, not the effect of how many compounds it produced or shared with its ancestor.

##### 3.1 Determine the number of compounds for each species and for each ancestral node.

We simply fixed the number of compounds produced by each species so that it was the same in both the null and real data for every species. We then tested for phylogenetic signal in the number of compounds produced by a species and used ancestral state reconstruction to estimate the number of compounds at each internal node in the phylogeny.

```
sampsByCompounds_pres_abs <- (sampsByCompounds > 0) * 1
ncomps <- rowSums(sampsByCompounds_pres_abs > 0)
ncomps <- setNames(ncomps, rownames(sampsByCompounds_pres_abs))
obj <- contMap(inga.tree_rooted, ncomps, plot = F)
plot(obj)
```

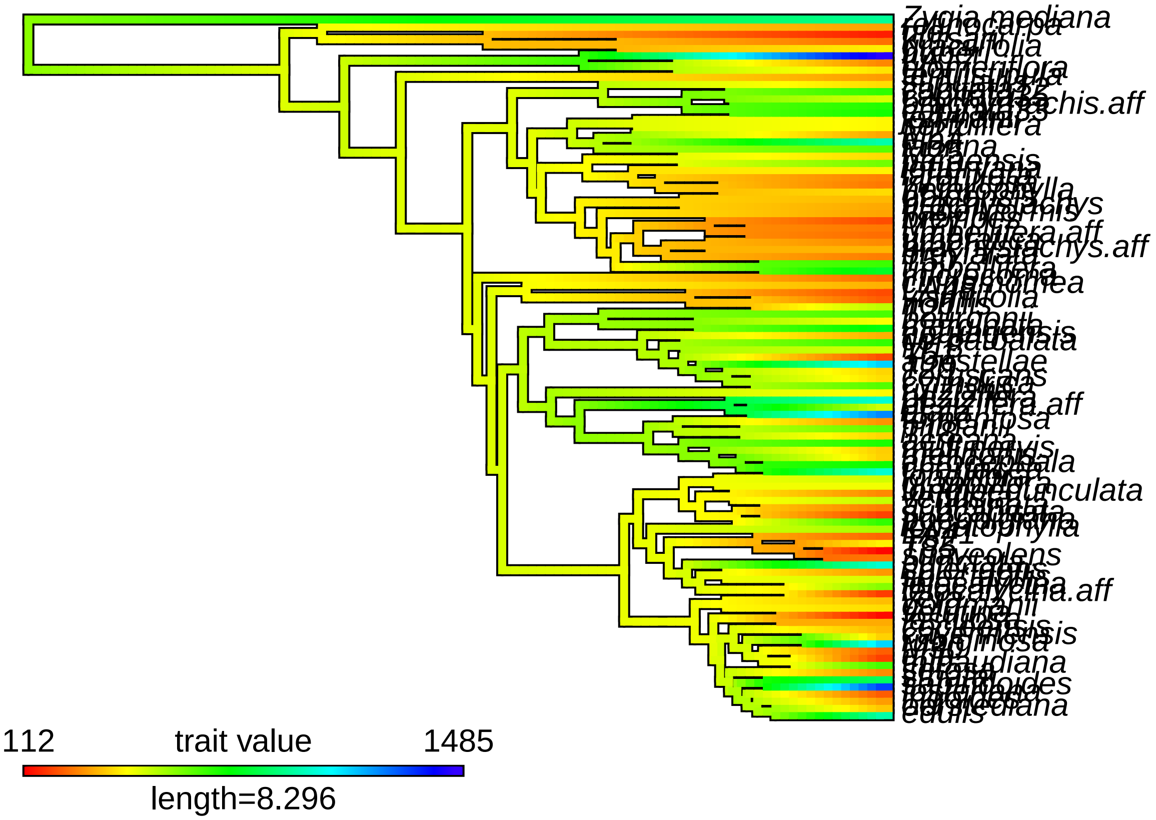

```
phylosig(inga.tree_rooted, ncomps, test = TRUE)
```

```
##
## Phylogenetic signal K : 0.404715
## P-value (based on 1000 randomizations) : 0.273
```

No significant phylogentic signal in the number of compounds shared.

For internal nodes we estimated ncompounds using ancestral state reconcstruction.

```
names_to_use <- row.names(read.csv(here("./results/ancestral_sampsBycompounds//
Ancestral_State_V1_all_compounds.
    row.names = 1))

fit <- fastAnc(inga.tree_rooted, ncomps, vars = TRUE, CI = TRUE)

anc_state_ncomps <- as.numeric(c(as.numeric(fit$ace), as.numeric(ncomps)))
anc_state_ncomps <- setNames(anc_state_ncomps, names_to_use)

plot(anc_state_ncomps, col = grepl("Node", names_to_use) + 1)
```

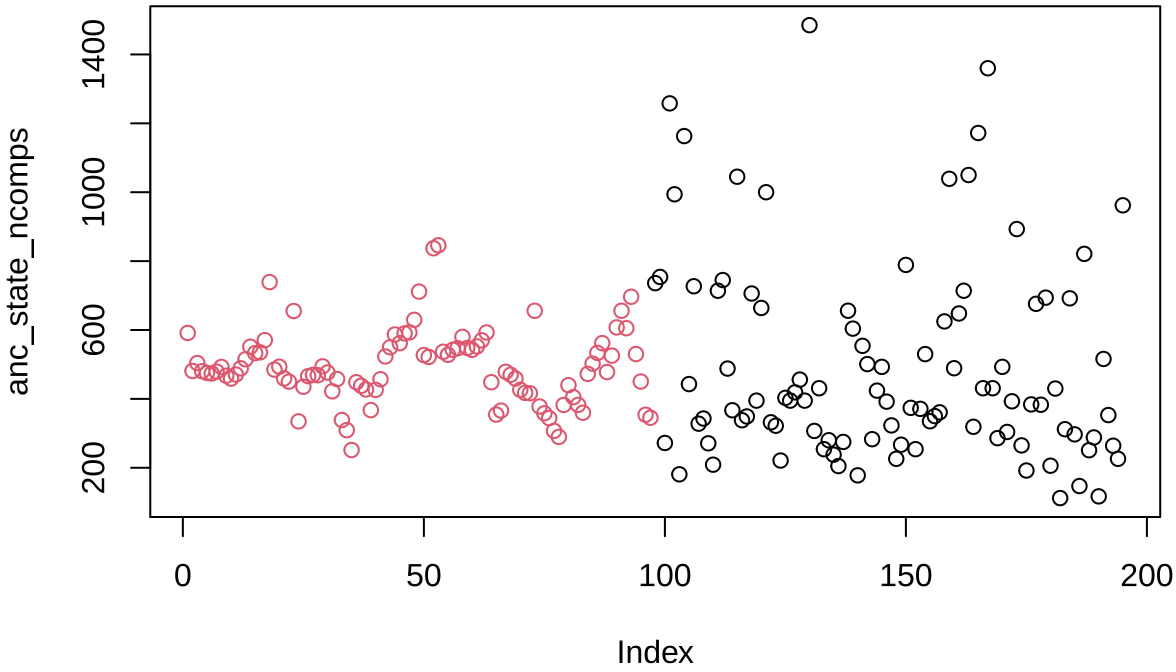

We then used these estimates to determine the number of compounds at each internal node in the null model.

##### 3.2 Determine the number of compounds shared between each species and its parent node.

To determine how many compounds were shared between a given species and a parent node we used ancestral state reconstruction to generate ancestral state (present/absent) of each compound at each internal node. From this ancestral SamplesByCompound maxtrix we determined the number of compounds shared between a species (phylogenetic tip) and its parent node.

The following code was used to generate a Node x compound matrix with present absence matrix estimated via ancestral state reconstruction. Note this code takes a long time to run, so I've saved the output in: ./results/

```
# library(parallel) library(doParallel) library(RMySQL) comp_rates <-
# data.frame() sampsByCompounds_pres_abs[1:10,1:10] com_comps <-
# sampsByCompounds_pres_abs[,colSums(sampsByCompounds_pres_abs)>1] #5578
# compounds dim(com_comps) com_comps <- data.frame(com_comps) names(com_comps) <-
# gsub('X','',names(com_comps)) com_comps[1:10,1:10] comps_list <-
# names(com_comps) all_comps <- colnames(sampsByCompounds_pres_abs) #comps_fail
# <- c('13','24') #comps_list <- comps_list[!comps_list %in% comps_fail] mydb =
# dbConnect(MySQL(), user= '',password = '!',dbname =
# 'inga_2015_06_01',host='mysql.chpc.utah.edu') select <- paste('SELECT * FROM
# `Ancestral_State_V1`') processed <- as.data.frame(dbGetQuery(mydb,select))
# ancestral_sampsbycomps <- as.data.frame(t(processed[,2:197]))
# names(ancestral_sampsbycomps) <- ancestral_sampsbycomps['compound',]
# ancestral_sampsbycomps <- ancestral_sampsbycomps[-1,] n_internal_nodes <-
# length(which(grepl('Node',row.names(ancestral_sampsbycomps))))
# length(comps_list) comps_list <- comps_list[!comps_list %in%
# as.character(processed$compound)] length(comps_list) #for all compounds in comp
# list do ancestral state reconstruction and upload to db #note this list
# excludes compounds that fail (appears to be ones where they are only present
# twice and in all cases they appear to not be sister species in the phylogeny
# but separated) cores <- detectCores() cl <- parallel::makeCluster(24,
```

```r
# setup_strategy = 'sequential') clusterEvalQ(cl,{ library(RMySQL) mydb =
# dbConnect(MySQL(), user= '',password = '!',dbname =
# 'inga_2015_06_01',host='mysql.chpc.utah.edu') NULL }) registerDoParallel(cl) #
# foreach(i = 1:length(comps_list) ,
# .packages=(c('phytools','caper')),.errorhandling = 'pass',.verbose = F) %dopar%
# { #if(i %% 100 == 0){print(paste('working on compound ', i, ' out of ',
# length(comps_list)))} ind_comp <- com_comps[,which(names(com_comps) ==
# comps_list[i])] ind_comp <-
# setNames(ind_comp,rownames(sampsByCompounds_pres_abs))
# obj<-contMap(inga.tree_rooted,ind_comp,plot=F) plot(obj) try({ mtrees <-
# make.simmap(inga.tree_rooted,ind_comp,nsim=100,model = 'ER',message = F)
# comp_rates_ind <- data.frame(compound = comps_list[i],npres = sum(ind_comp),
# tot_trans = colMeans(summary(mtrees)$count)[1], trans_0_to_1 =
# colMeans(summary(mtrees)$count)[2], trans_1_to_0 =
# colMeans(summary(mtrees)$count)[3]) lambda_phyl <-
# phylosig(inga.tree_rooted,ind_comp,test=TRUE,method = 'lambda')
# comp_rates_ind$phyl_sig_lambda <- lambda_phyl$lambda
# comp_rates_ind$phyl_sig_lambda_pval <- lambda_phyl$P BlomK_phyl <-
# phylosig(inga.tree_rooted,ind_comp,test=TRUE) comp_rates_ind$phyl_sig_K <-
# BlomK_phyl$K comp_rates_ind$phyl_sig_K_pval <- BlomK_phyl$P ind_comp_df <-
# as.data.frame(ind_comp) ind_comp_df$species <- row.names(ind_comp_df) D_phyl <-
# phylo.d(ind_comp_df, inga.tree_rooted,names.col = species,binvar = ind_comp,
# permut = 1000) comp_rates_ind$phyl_sig_D <- D_phyl$DEstimate
# comp_rates_ind$phyl_sig_D_pval_rand <- D_phyl$Pval0
# comp_rates_ind$phyl_sig_D_pval_brown <- D_phyl$Pval1
# dbWriteTable(mydb,'Ancestral_State_comp_rates',comp_rates_ind,
# field.types=names(comp_rates_ind),row.names=F,overwrite=F, append=T) pd <-
# summary(mtrees) anc_state <- data.frame(anc_state =
# as.numeric(c(round(pd$ace[,2],0),ind_comp))) row.names(anc_state)=
# c(row.names(pd$ace),row.names(sampsByCompounds_pres_abs)) names(anc_state) =
# comps_list[i] #return(anc_state) anc_state_1 <- cbind(data.frame(compound =
# as.numeric(names(anc_state))),t(anc_state)) names(anc_state_1)[2:98] <-
# paste('Node_',names(anc_state_1)[2:98],sep='')
# dbWriteTable(mydb,'Ancestral_State_V1',anc_state_1,
# field.types=names(anc_state_1),row.names=F,overwrite=F, append=T) },silent = T)
# } #now go back and update the list to deterimine qhich compounsd are missing
# mydb = dbConnect(MySQL(), user= '',password = '!',dbname =
# 'inga_2015_06_01',host='mysql.chpc.utah.edu') select <- paste('SELECT * FROM
# `Ancestral_State_V1`') processed <- as.data.frame(dbGetQuery(mydb,select))
# ancestral_sampsbycomps <- as.data.frame(t(processed[,2:197]))
# names(ancestral_sampsbycomps) <- ancestral_sampsbycomps['compound',]
# ancestral_sampsbycomps <- ancestral_sampsbycomps[-1,] n_internal_nodes <-
# length(which(grepl('Node',row.names(ancestral_sampsbycomps))))
# length(comps_list) missing_comps <- all_comps[!all_comps %in%
# as.character(processed$compound)] for (i in 1:length(missing_comps)){ ind_comp
# <- sampsByCompounds_pres_abs[,which(colnames(sampsByCompounds_pres_abs) ==
# missing_comps[i])] ind_comp <-
# setNames(ind_comp,rownames(sampsByCompounds_pres_abs))
# #obj<-contMap(inga.tree_rooted,ind_comp,plot=F) #plot(obj) anc_state <-
# data.frame(anc_state = as.numeric(c(rep(0,n_internal_nodes),ind_comp)))
# row.names(anc_state)= c(row.names(pd$ace),row.names(sampsByCompounds_pres_abs))
# names(anc_state) = missing_comps[i] #return(anc_state) anc_state_1 <-
# cbind(data.frame(compound = as.numeric(names(anc_state))),t(anc_state))
```

```r
# names(anc_state_1)[2:98] <- paste('Node_',names(anc_state_1)[2:98],sep='')
# dbWriteTable(mydb,'Ancestral_State_V1',anc_state_1,
# field.types=names(anc_state_1),row.names=F,overwrite=F, append=T)}

# finally combine all of these into a single data.frame that should contain all
# compounds

# mydb = dbConnect(MySQL(), user= '*****',password = '***',dbname =
# 'inga_2015_06_01',host='mysql.chpc.utah.edu') select <- paste('SELECT * FROM
# `Ancestral_State_V1`') processed <- as.data.frame(dbGetQuery(mydb,select))
# ancestral_sampsbycomps <- as.data.frame(t(processed[,2:197]))
# names(ancestral_sampsbycomps) <- ancestral_sampsbycomps['compound',]
# ancestral_sampsbycomps <- ancestral_sampsbycomps[-1,] ancestral_sampsbycomps <-
# ancestral_sampsbycomps[,order(as.numeric(colnames(ancestral_sampsbycomps)))]
# dim(ancestral_sampsbycomps)
# write.csv(ancestral_sampsbycomps,here('./results/ancestral_sampsBycompounds//
Ancestral_State_V1_all_compounds.

ancestral_sampsbycomps <- read.csv(here("./results/ancestral_sampsBycompounds//
Ancestral_State_V1_all_compounds.
    row.names = 1)
names(ancestral_sampsbycomps) <- gsub("X", "", names(ancestral_sampsbycomps))
```

```r
ancestral_sampsbycomps[1:10, 1:10]
```

```
##          2 3 5 6 7 8 9 11 12 13
## Node_99  0 0 0 0 0 0 0  0  0  0
## Node_100 0 0 0 0 0 0 0  0  0  0
## Node_101 0 0 0 0 0 0 0  0  0  0
## Node_102 0 0 0 0 0 0 0  0  0  0
## Node_103 0 0 0 0 0 0 0  0  0  0
```

Finally, from the above estimates we generated a data.frame in which we calculated the following for each parent and child in the phylogeny: a) ncomps_ancestral: estimated number of compounds in parent node b) ncomps_shared: number of compounds shared between the parent and child node c) ncomps_lost: number of compounds lost (ncomps_ancestral - ncomps_shared)

```r
ncomps_expected_vs_reconstructed_recent_nodes <- data.frame(node =
row.names(ancestral_sampsbycomps)[grepl("Node",
    row.names(ancestral_sampsbycomps))], expected =
    as.numeric(anc_state_ncomps[grepl("Node",
    row.names(ancestral_sampsbycomps))]), reconstructed =
    as.numeric(rowSums(ancestral_sampsbycomps > 0)[grepl("Node",
    row.names(ancestral_sampsbycomps))]))

ncomps_expected_vs_reconstructed <- ncomps_expected_vs_reconstructed_recent_nodes[,
    2:3]
row.names(ncomps_expected_vs_reconstructed) <- ncomps_expected_vs_reconstructed_recent_nodes$node

tree <- inga.tree_rooted
total_nodes <- length(tree$tip.label) + Nnode(tree)
cur_node <- 1 + length(tree$tip.label)
```

```r
nodes_to_do <- data.frame()
for (node in 99:195) {
    next_nodes <- Descendants(tree, node, "children")
    if (length(next_nodes) > 0) {
        nodes_to_do <- rbind(nodes_to_do, data.frame(parent = node, child = next_nodes))
    }
}

recent_nodes <- nodes_to_do[nodes_to_do$child < 99, ]
recent_nodes$child_names <- tree$tip.label[recent_nodes$child]
recent_nodes$parent_names <- paste("Node_", recent_nodes$parent, sep = "")

recent_nodes_1 <- merge(recent_nodes, ncomps_expected_vs_reconstructed, by.x = "parent_names",
    by.y = 0)

# recent_nodes_1$child_names[which(recent_nodes_1$child_names ==
# 'leiocalycina.aff')] <- 'leiocalycina.af'
# recent_nodes_1$child_names[which(recent_nodes_1$child_names == 'tomentosa')] <-
# 'omentosa'

transitions <- data.frame()
for (row in 1:nrow(recent_nodes_1)) {
    comp_pool <- ancestral_sampsbycomps[c(recent_nodes_1$parent_names[row], recent_nodes_1
    $child_names[row]),
        ]
    ncomps_ancestral <- recent_nodes_1$expected[row]
    ncomps_shared <- sum(colSums(comp_pool) == 2)
    ncomps_new <- sum(as.numeric(comp_pool[2, ]) == 1 & as.numeric(comp_pool[1, ]) ==
        0)
    transitions <- rbind(transitions, data.frame(parent = recent_nodes_1$parent_names[row],

        child = recent_nodes_1$child_names[row], ncomps_ancestral = ncomps_ancestral,
        ncomps_shared = ncomps_shared, ncomps_new = ncomps_new, ncomps_lost = ncomps_ancestral -
            ncomps_shared))
}

transitions$ncomps_child <- transitions$ncomps_shared + transitions$ncomps_new
transitions$per_shared_child <- transitions$ncomps_shared/transitions$ncomps_child
```

#### 4.0 Putting this all together into our complete null model

Below is the code we used to generate 98 iterations of the above null model. Each iteration was seeded with a different randomly generated chemical profile.

Null models used in this paper are saved in ./results/null_model/1_SampsByComps_Null_Model

```r
# nodes_to_do <- data.frame() for(node in 99:195){ next_nodes <-
# Descendants(tree,node,'children') if(length(next_nodes) >0) {nodes_to_do <-
# rbind(nodes_to_do,data.frame(parent = node,child = next_nodes))} }
# nodes_to_do$child_names <- nodes_to_do$child
# nodes_to_do$child_names[nodes_to_do$child < 99] <-
# tree$tip.label[nodes_to_do$child[nodes_to_do$child < 99]]
# nodes_to_do$child_names[nodes_to_do$child > 99] <-
```

```
# paste('Node_',nodes_to_do$child[nodes_to_do$child > 99],sep='')
# nodes_to_do$parent_names <- paste('Node_',nodes_to_do$parent,sep='')
# comp_frequency <- colSums(sampsByCompounds>0) cl <- parallel::makeCluster(40,
# setup_strategy = 'sequential') registerDoParallel(cl) foreach(i =
# 1:nrow(sampsByCompounds_rand),.packages = c('ape','phangorn','here')) %dopar% {
# #random_seed <- sample(1:nrow(sampsByCompounds_rand),1) starting_node <-
# sampsByCompounds_rand[i,] sampsByCompounds_Evolved <- data.frame()
# sampsByCompounds_Evolved <-rbind(sampsByCompounds_Evolved,starting_node)
# sampsByCompounds_Evolved$node_label <- 'Node_99' sampsByCompounds_Evolved <-
# sampsByCompounds_Evolved[,c(ncol(sampsByCompounds_Evolved),2:ncol(sampsByCompounds_Evolved)-1)]
# names(sampsByCompounds_Evolved) <- c('node_label',names(sampsByCompounds))
# sampsByCompounds_Evolved[,1:10] for(n in 1:nrow(nodes_to_do)){ child =
# as.character(nodes_to_do$child_name[n]) parent =
# as.character(nodes_to_do$parent_names[n]) evolved_abund_vect <-
# inherit_comps(child = child,parent = parent,starting_node = starting_node)
# evolved_abund_vect <- sample_new_comps(parent = parent,child =
# child,evolved_abund_vect = evolved_abund_vect) sampsByCompounds_Evolved <-
# rbind(sampsByCompounds_Evolved,c(child,evolved_abund_vect)) }
# write.csv(sampsByCompounds_Evolved,here(paste('./results/null_model/1_SampsByComps_Null_Model/Evolved_
# = F) }
```
